# Mechanistic insights into direct DNA and RNA strand transfer and dynamic protein exchange of SSB and RPA

**DOI:** 10.1101/2025.04.01.643995

**Authors:** Tapas Paul, I-Ren Lee, Sushil Pangeni, Fahad Rashid, Olivia Yang, Edwin Antony, James M. Berger, Sua Myong, Taekjip Ha

## Abstract

Single-stranded DNA-binding proteins (SSBs) are essential for genome stability, facilitating replication, repair, and recombination by binding ssDNA, recruiting other proteins, and dynamically relocating in response to cellular demands. Using single-molecule fluorescence resonance energy transfer (smFRET) assays, we elucidated the mechanisms underlying direct strand transfer from one locale to another, protein exchange, and RNA interactions at high resolution. Both bacterial SSB and eukaryotic replication protein A (RPA) exhibited direct strand transfer to competing ssDNA, with rates strongly influenced by ssDNA length. Strand transfer proceeded through multiple failed attempts before a successful transfer, forming a ternary intermediate complex with transient interactions, supporting a direct transfer mechanism. Both proteins efficiently exchanged DNA-bound counterparts with freely diffusing molecules, while hetero-protein exchange revealed that SSB and RPA could replace each other on ssDNA in a length-dependent manner, indicating that protein exchange does not require specific protein-protein interactions. Additionally, both proteins bound RNA and underwent strand transfer to competing RNA, with RPA demonstrating faster RNA transfer kinetics. Competitive binding assays confirmed a strong preference for DNA over RNA. These findings provide critical insights into the dynamic behavior of SSB and RPA in nucleic acid interactions, advancing our understanding of their essential roles in genome stability, regulating RNA metabolism, and orchestrating nucleic acid processes.

## INTRODUCTION

Single-stranded DNA-binding proteins (SSBs) such as bacterial SSB and eukaryotic replication protein A (RPA) are indispensable for preserving genome stability by orchestrating DNA replication, repair, and recombination (1,2). These proteins bind single-stranded DNA (ssDNA) with high affinity (dissociation constant, K_D_ ~ 10^−10^ M) and prevent secondary structure formation while stabilizing intermediates required for essential cellular processes (1,3–5). Despite differences in structural composition, SSB in bacteria and RPA in eukaryotes and archaea exhibit functional similarities (6). SSB binds ssDNA cooperatively, wrapping the strand in a protective protein coat, whereas RPA employs its modular DNA-binding domains in a flexible and dynamic manner, allowing adaptation to varying DNA contexts (7–10). Both proteins play critical roles in replication fork stability, particularly in lagging-strand synthesis, where their redistribution facilitates DNA polymerase progression and ensures timely repair of damaged regions (1,11,12).

Strand transfer and protein exchange are fundamental to the efficient function of SSB and RPA in dynamic cellular environments. During DNA replication and repair, the ability of these proteins to rapidly transfer from one ssDNA segment to another (‘strand transfer’) or exchange with freely diffusing proteins (‘protein exchange’) ensures that critical processes proceed unimpeded (1,12–14). This adaptability is essential for replacing DNA-bound proteins during transitions in DNA metabolic pathways and recruiting repair or recombination factors. Strand transfer, in particular, is crucial for maintaining the continuity of replication forks and resolving obstacles such as nucleic acid secondary structure or damaged templates (11,14–16). The kinetics of strand transfer and protein exchange provide critical insights into these mechanisms, revealing the rate-limiting steps and intermediate states that govern these processes (17–21).

SSB and RPA exhibit distinct diffusion properties integral to their functions. SSB diffuses along ssDNA with a diffusion coefficient of ~270 nt^2^/s on short ssDNA, which increases significantly (~170,000 nt^2^/s) for long ssDNA under low tension (22–25). In comparison, RPA demonstrates a higher diffusion coefficient (~3,000 nt^2^/s) on short ssDNA, attributed to its ability to dynamically rearrange its DNA-binding domains during movement, which increases up to ~20,000 nt^2^/s on long ssDNA (24,26). Mechanisms such as hopping and intersegmental transfer allow both proteins to traverse non-contiguous ssDNA regions, supporting efficient redistribution during replication and repair (9,23,27). Furthermore, the salt-dependent binding modes of SSB (e.g., transitioning between 35-nt and 65-nt ssDNA binding modes) and the configurational adaptability of RPA enable these proteins to fine-tune their interactions with DNA, facilitating strand transfer and protein exchange (7,9,28).

Strand transfer and protein exchange are central to SSB and RPA functionality, allowing homotypic and heterotypic exchanges. These processes are crucial for the replacement of DNA-bound proteins during transitions in metabolic pathways. For example, direct strand transfer occurs via a ternary intermediate state where both proteins bind competing ssDNA strands simultaneously, enabling efficient displacement (12,29–31). Hetero-protein exchange dynamics highlight their cooperative roles in recruiting repair and recombination factors, while their ability to bind RNA adds a layer of versatility (32,33). Interestingly, while SSB and RPA are well-studied in the context of DNA metabolism, their interactions with RNA remain less explored. SSBs can have a similar binding affinity for RNA and DNA (34,35) and can prevent the formation of a DNA-RNA hybrid (36,37), yet the dynamics and functional significance of RNA binding remains largely unexplored. RPA also interacts with RNA during processes such as transcription-coupled repair, highlighting its potential roles in RNA metabolism and cellular regulation (38–40). Exploring these interactions could reveal novel dimensions of their functionality, particularly in pathways where DNA and RNA substrates overlap or compete for binding.

Despite these advances, the mechanistic underpinnings of SSB and RPA strand transfer, protein exchange, and RNA binding remain incomplete. Here, we employ single-molecule FRET to expand upon previous research by investigating the direct strand transfer and protein exchange dynamics of SSB and RPA on ssDNA and RNA. Our study addresses key questions regarding how these proteins undergo strand transfer between ssDNA molecules of varying lengths and their ability to exchange with each other on ssDNA. Additionally, we examine the binding competition between DNA and RNA for both proteins, providing new insights into their roles in genome stability and RNA metabolism. By combining kinetic measurements with mechanistic insights, this study offers a comprehensive view of SSB and RPA functionality, with implications for understanding nucleic acid processing in various cellular contexts.

## MATERIALS AND METHODS

### Preparation of DNA constructs

The HPLC-purified DNA and RNA oligonucleotides (listed in Supplementary Table S1) containing biotin for immobilization and either Cy3, Cy5, or amine modifications were procured from IDT. Amine-modified oligonucleotides were labeled with NHS ester-conjugated fluorescent dyes following established protocols (41). Partial duplex DNA constructs (10 µM) were prepared by mixing a biotin-conjugated DNA strand with its complementary strand at a molar ratio of 1:1.2 (biotinylated:non-biotinylated). DNA strands were annealed in T50 buffer (10 mM Tris-HCl, pH 7.5, and 50 mM NaCl) using a thermocycler. The mixture was heated to 95°C for 2 minutes, then gradually cooled at a rate of 2°C/min until 40°C, followed by cooling at 5°C/min until 4°C (42). The annealed constructs were stored at −20°C and freshly re-annealed before use. All buffers were prepared using Milli-Q water and filtered through 0.22 µm membrane filters.

### Protein purification and labelling

SSB protein was cloned into the pRSF vector, and a G26C mutation was introduced for labeling purposes. To express SSB, the pRSF-SSB G26C vector was transformed into *E. coli* BL21(DE3) cells. The cells were grown at 37°C and induced with 0.5 mM IPTG when the optical density (OD) at 600 nm reached 0.8. They were harvested after an additional 3-hour incubation. Cells were resuspended in lysis buffer containing 50 mM HEPES (pH 7.5), 200 mM NaCl, 15% glycerol, 2 mM DTT, and a protease inhibitor cocktail. To purify SSB, the cells were thawed and lysed by adding lysozyme (2 mg/mL), incubating on ice for 30 minutes, followed by sonication. The lysate was clarified by centrifugation. SSB was precipitated by adding polyethylenimine (PEI) to a final concentration of 0.2%. The pellet was then solubilized in buffer containing 50 mM HEPES (pH 7.5), 500 mM NaCl, 15% glycerol, and 2 mM DTT. SSB was further precipitated by adding ammonium sulfate to a final concentration of 30%. The pellet was solubilized in buffer containing 50 mM HEPES (pH 7.5), 50 mM NaCl, 15% glycerol, and 1 mM TCEP. The conductivity of the sample was adjusted to 100 mM NaCl before loading onto a HiTrap Heparin column. SSB was eluted using a linear gradient of 100-1000 mM NaCl in the same buffer.

For labeling, a 10-fold molar excess of Cy3-maleimide dye was added, and the reaction was incubated overnight on ice. The reaction was quenched by adding an excess of 1 mM DTT. Free unreacted dye was removed using PD-10 desalting columns. Labeled SSB was further purified using a HiLoad Superdex-200 column pre-equilibrated with buffer containing 50 mM HEPES (pH 7.5), 500 mM NaCl, 15% glycerol, and 1 mM TCEP. Peak fractions were collected, concentrated, aliquoted, and stored at −80°C. Labeling efficiency was estimated to be above 90% based on absorbance measurements.

RPA protein was purified and labeled (with MB543) as described (9,43).

### Single-molecule FRET assays

Single-molecule FRET experiments were performed on a custom-built prism-type total internal reflection (PTIR) inverted fluorescence microscope (Olympus IX 71) as previously described (44,45). Experiments were conducted on quartz slides coated with polyethylene glycol (PEG) to prevent non-specific interactions between excess DNA, RNA, and protein. Quartz slides and glass coverslips were pre-drilled, washed thoroughly with methanol and acetone, and sonicated in 1 M potassium hydroxide. Slides were burned for 2-3 minutes, while coverslips were sterilized by brief flaming (4-5 passes). Both slides and coverslips were then treated with aminosilane for 30 minutes and coated with a mixture of 98% mPEG (mPEG-5000; Laysan Bio) and 2% biotin-PEG (biotinPEG-5000; Laysan Bio) (46).

A microfluidic sample chamber was created by assembling the PEG/biotin-PEG coated slide and coverslip. Stocks of annealed partial duplex DNA or RNA (10 µM) were diluted to 15-20 pM and immobilized on the PEG-passivated surface via biotin-NeutrAvidin (50 µg/ml) interactions. Unbound molecules were removed by washing with excess buffer. smFRET measurements were performed in imaging buffer containing 10 mM Tris-HCl (pH 7.5), 100 mM or 300 mM NaCl, 10% glycerol, and an oxygen scavenging system (10 mM Trolox, 0.5% glucose, 1 mg/ml glucose oxidase, and 4 µg/ml catalase) to minimize photobleaching and improve dye stability. All the smFRET measurements were carried out at room temperature (~23°C ± 2°C). All smFRET measurements were repeated either in the same channel by following the regeneration methods or in a new channel on different days (46).

### smFRET data acquisition and analysis

A solid-state diode laser (532 or 634 nm; Compass 315M, Coherent) was used to generate an evanescent field through PTIR to excite the fluorophores (Cy3 or Cy5). Fluorescence emissions from Cy3 (donor) and Cy5 (acceptor) were collected using a water immersion objective and projected onto an EMCCD camera (Andor) via a dichroic mirror (cutoff = 630 nm). Data were recorded with a 50 ms frame integration time, processed with an IDL scripts (http://www.exelisvis.co.uk/ProductsServices/IDL.aspx) and analyzed using MATLAB scripts (https://www.mathworks.com/). The FRET efficiency (*E*_*FRET*_) was calculated using *I*_*A*_/(*I*_*D*_ + *I*_*A*_), where *I*_*D*_ and *I*_*A*_ represent the intensity of donor and acceptor respectively. FRET histograms were generated from >4,000 molecules (21 frames from 20 short movies) across different imaging surfaces. Alternating green and red laser excitation (10 frames each, separated by a dark frame) excluded donor-only molecules in low *E*_*FRET*_ regions. Donor leakage was corrected based on donor-only *E*_*FRET*_ values. Histograms were normalized and fitted with multi-peak Gaussian distributions.

### Strand transfer and protein exchange

Strand transfer assays were performed using microfluidic imaging chambers prepared as described above (45,46). SSB or RPA proteins were first bound to immobilized DNA or RNA constructs, followed by the addition of competing ssDNA or RNA at varying concentrations. FRET histograms were fitted to Gaussian distributions, and the bound protein fraction was calculated over time to determine binding kinetics. For real-time smFRET measurements, proteins suspended in imaging buffer were introduced into the chamber via a syringe pump at a flow rate of 20 µl/s. Cy3-labeled ssDNA was added to visualize intermediate complex formation during strand transfer in real time.

Protein exchange experiments were conducted similarly to that of strand exchange. Cy5-labeled partial duplex DNA constructs were immobilized to ensure optimal molecule binding on the surface (~300). Labeled SSB or RPA was added, followed by unlabeled protein at varying concentrations. Fluorescence signals decreased over time as labeled proteins exchanged with unlabeled counterparts. Short movies (2 s) were recorded at different time intervals, and spot counts were plotted over time. Exchange kinetics were calculated using single-exponential decay fitting in Origin. Hetero-protein exchange was measured similarly by introducing the corresponding proteins into the reservoir and monitoring *E*_*FRET*_ changes during the transition. Kinetics were calculated from flow initiation to complete exchange, based on FRET efficiency changes. More than 200 single-molecule time traces were analyzed for each condition.

## RESULTS

### SSB binding and direct transfer kinetics on ssDNA

We developed a single-molecule FRET assay to investigate the direct transfer of bound SSB between single-stranded DNAs (Figure 1A). In this assay, we used Cy3 (donor)- and Cy5 (acceptor)-labeled partial duplex (pd) DNA (18 bp) containing a poly-thymine stretch (dT_70_) at the 3′ end (pdT_70_), immobilized on a slide by a biotin-NeutrAvidin linkage. Upon addition of SSB (10 nM) to pdT_70_ in a buffer containing 300 mM NaCl, the low-*E*_*FRET*_ peak (~0.2) corresponding to DNA-only shifted to a high-*E*_*FRET*_ peak (~0.8), indicating complete SSB binding (Figure 1B). The high-*E*_*FRET*_ is consistent with previous findings that the DNA wraps around the protein in its (SSB)_65_ binding mode in high salt conditions (22,28). The bound SSB remained stable for hours even after excess unbound SSB proteins were removed from the solution.

**Figure 1.**
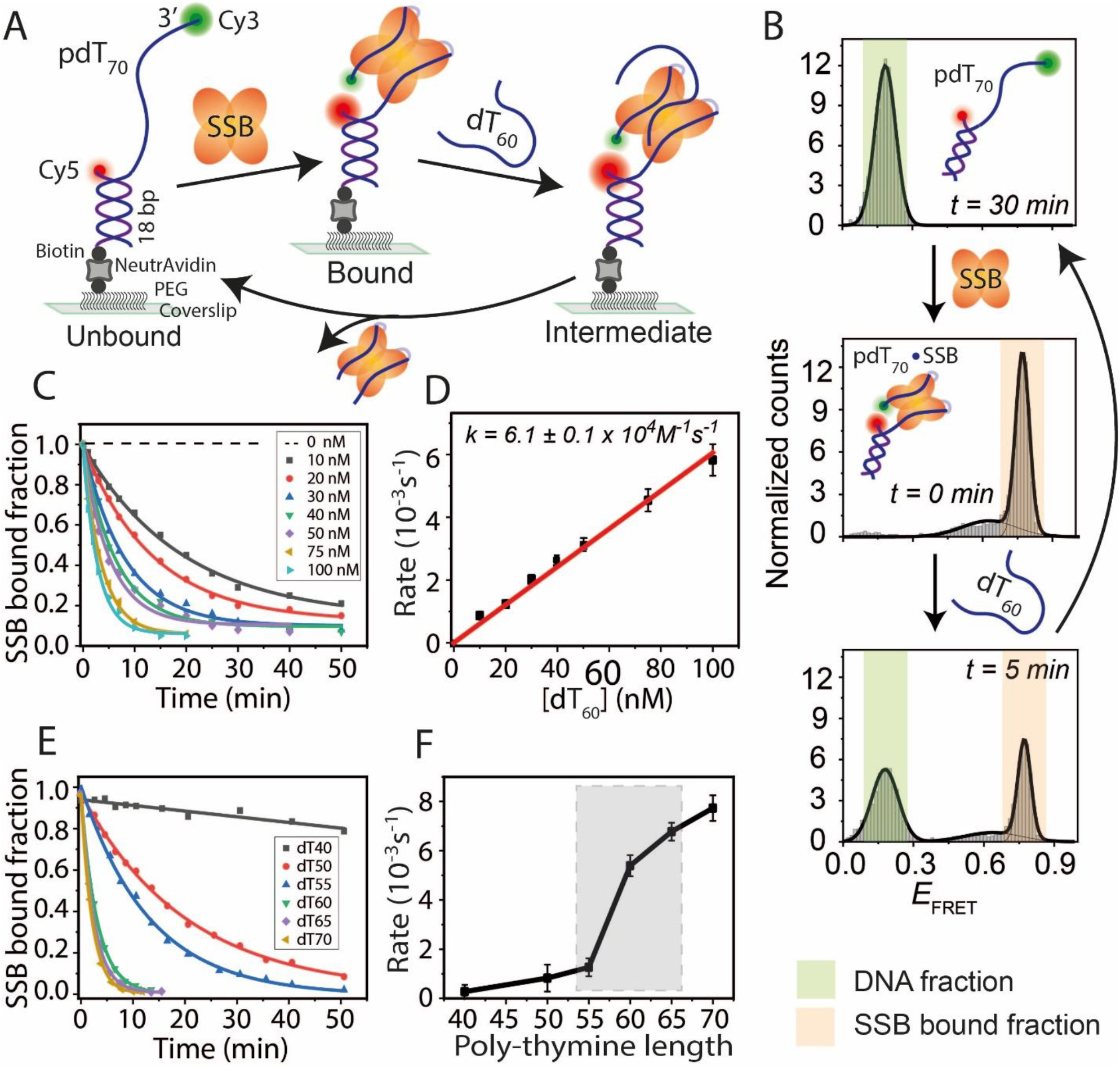
SSB binding and length dependent direct transfer kinetics on ssDNA. (A) Schematic of smFRET constructs showing a partial DNA duplex with a 70-nt poly-thymine overhang (pdT_70_). Sequential addition of SSB binds to the ssDNA overhang, and competing free ssDNA (dT_60_) facilitates strand transfer to regenerate the tethered DNA. (B) FRET histograms of pdT70 before (top, light green), after SSB binding (middle, light orange), and during SSB transfer (bottom) upon dT_60_ addition at the indicated time points. Time, t=30 min (top histogram) represents the complete strand transfer of bound SSB. (C) Single-exponential fitting of the SSB-bound fraction at varying concentrations of competing dT_60_. (D) Linear fit of SSB transfer rates at different dT_60_ concentrations. (E) Single-exponential fitting of SSB-bound fractions with competing ssDNA of different lengths (dT_40_ to dT_70_ at 100 nM). (F) SSB transfer rates plotted against competing ssDNA length, with a significant rate jump highlighted by the shaded gray region.

To assess SSB transfer between DNA strands, we introduced competing ssDNA (60-nt poly-thymine, dT_60_) into the sample chamber containing immobilized SSB-bound complexes. We hypothesized that the bound protein would transfer to the competing strand, regenerating pdT_70_ without a bound SSB. Over time, the fraction of surface-tethered DNA bound by SSB, represented by the high-*E*_*FRET*_ population, decreased, while the low-*E*_*FRET*_ DNA-only population increased (Figure 1B). To evaluate the efficiency of competing ssDNA-dependent SSB transfer, we titrated dT_60_ from 10 nM to 100 nM. Kinetic analysis was performed by plotting the SSB-bound fraction as a function of time. At all dT_60_ concentrations tested, the data fit well with single-exponential decay curves (Figure 1C), consistent with pseudo-first-order kinetics as observed in previous bulk solution kinetics studies (29). The transfer rates exhibited a linear dependence on dT_60_ concentration (Figure 1D), suggesting that one molecule of competing ssDNA was involved in the rate-determining step of SSB transfer. This finding supports a direct transfer mechanism rather than a cooperative mechanism, which would involve more than one competing ssDNA molecule in the rate-determining step (29). A similar mechanism was observed with a shorter DNA (40-nt of poly-thymine, pdT_40_) bound by SSB and transferring the bound SSB to a competing dT_40_ ssDNA in a buffer containing 100 mM NaCl (Supplementary Figure S1). The apparent SSB transfer rate for pdT_40_ to dT_40_ (9.8 ± 0.2 × 10^4^ M^−1^ s^−1^) was similar to that of pdT_70_ to dT_60_ transfer (6.1 ± 0.1 × 10^4^ M^−1^ s^−1^).

Next, to examine the effect of competing ssDNA length on SSB transfer, we varied the ssDNA length from dT_40_ to dT_70_ while keeping the same tethered pdT_70_-SSB complex. These lengths were chosen based on SSB’s known ssDNA binding modes (28,47). The overall transfer reaction was faster when longer competing ssDNA molecules were used (Figure 1E, F). Interestingly, a significant jump in the transfer rate was observed between dT_55_ and dT_60_. Altogether, the transfer of bound SSB to the competing ssDNA follows pseudo-first-order transfer kinetics and is ssDNA length-dependent.

### Binding Kinetics and direct transfer of RPA on ssDNA

Next, we investigated the direct transfer of bound *Saccharomyces cerevisiae*-RPA using the same single-molecule FRET assay (Figure 2A). The surface-tethered partial duplex (pd) DNA contained 40 nt of poly-thymine (pdT_40_) at the 3′ end. Upon addition of RPA (10 nM) to pdT_40_ in a buffer containing 100 mM NaCl, the low-*E*_*FRET*_ peak (~0.3) corresponding to DNA-only shifted to an even lower-*E*_*FRET*_ peak (~0.1), indicating complete RPA binding and stretching of the bound ssDNA (Figure 2B). This binding remained stable for hours, even after washing away excess unbound RPA from the solution. Under these conditions only one RNA can be bound to the ssDNA (48).

**Figure 2.**
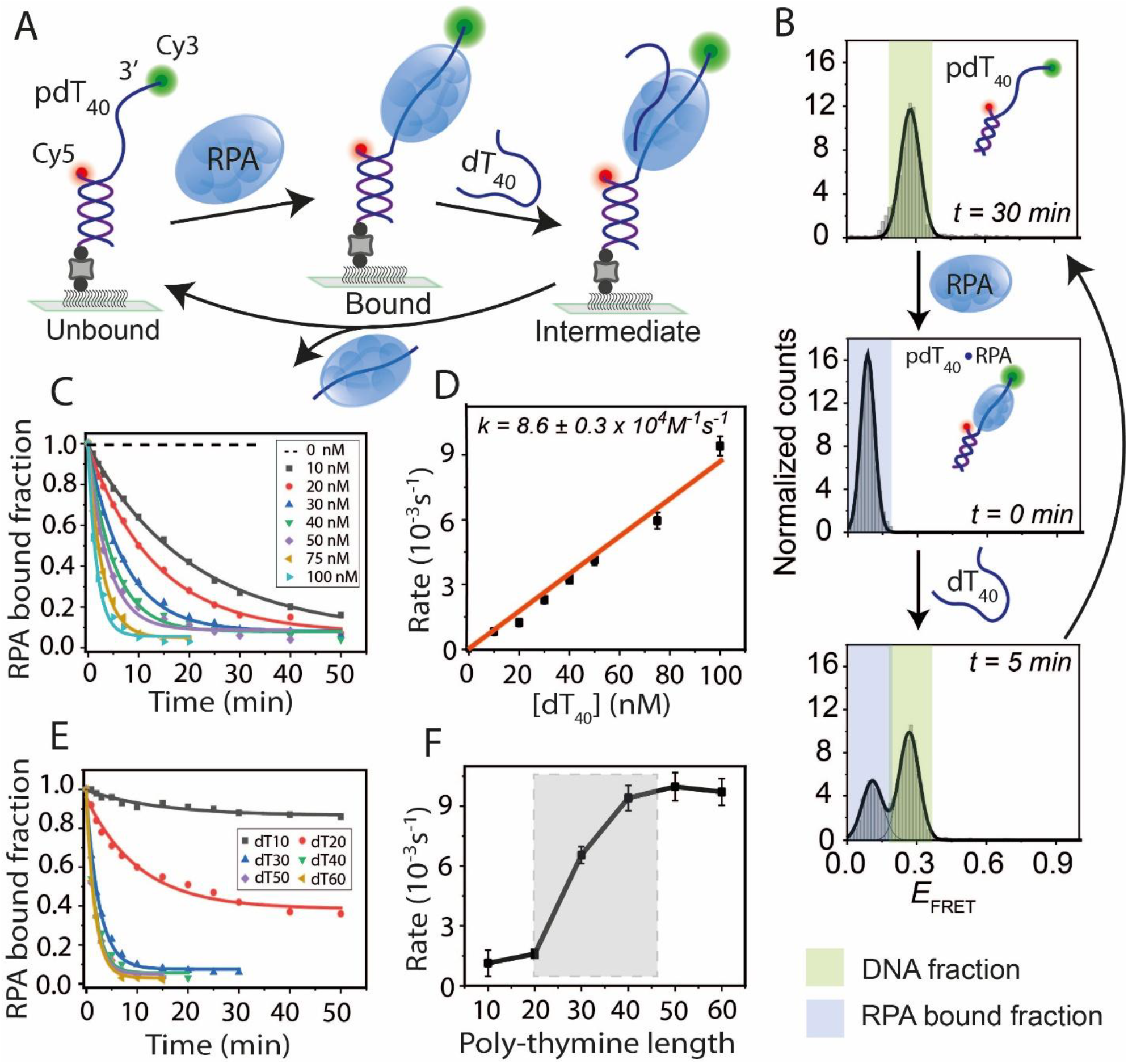
Length dependent binding kinetics and direct transfer of budding yeast RPA on ssDNA. (A) Schematic of smFRET constructs with a partial duplex containing a 40-nt poly-thymine overhang (pdT_40_). RPA binds to pdT_40_ and competing free ssDNA (dT_40_) induces strand transfer to regenerate the tethered DNA. (B) FRET histograms of pdT_40_ before (top, light green), after RPA binding (middle, light blue), and during RPA transfer (bottom) upon dT_40_ addition at indicated times. Time, t=30 min (top histogram) represents the complete strand transfer of bound RPA. (C) Single-exponential fitting of RPA-bound fractions at different dT_40_ concentrations. (D) Linear fit of RPA transfer rates at varying dT_40_ concentrations. (E) Single-exponential fitting of RPA-bound fractions with competing ssDNA of different lengths (dT_10_ to dT_60_ at 100 nM). (F) RPA transfer rates plotted against competing ssDNA length, with a significant rate jump highlighted by the shaded gray box.

To study RPA strand transfer, we introduced competing dT_40_ ssDNA into the tethered complex. Over time, the RPA-bound *E*_*FRET*_ population decreased, while the DNA-only *E*_*FRET*_ population increased (Figure 2B). Kinetic analysis was performed by titrating dT_40_ at various concentrations, and the RPA bound fraction was plotted as a function of time (Figure 2C). The data fit well with single-exponential decay curves, consistent with pseudo-first-order kinetics, as observed for SSB. The transfer rates exhibited a linear dependence on the competing ssDNA concentration (Figure 2D). A similar strand transfer experiment was performed using the pdT_70_-RPA complex with dT_60_ as the competing ssDNA under the same buffer conditions (Supplementary Figure S2). The apparent RPA transfer rates for both cases were nearly identical (8.6 ± 0.3 × 10^4^ M^−1^ s^−1^ for pdT_40_ to dT_40_ and 9.1 ± 0.4 × 10^4^ M^−1^ s^−1^ for pdT_70_ to dT_60_). Furthermore, a similar transfer reaction from pdT_40_ to dT_40_ was observed for human-RPA, with an apparent transfer rate (8.3 ± 0.6 × 10^4^ M^−1^ s^−1^) comparable to that of yeast-RPA (Supplementary Figure S3), suggesting that yeast- and human-RPA are nearly identical in their ability to transfer from one DNA strand to another.

To assess the effect of ssDNA length on RPA transfer, we varied the competing ssDNA length from dT_10_ to dT_60_ for the tethered pdT_40_. We found that longer ssDNA facilitated faster transfer (Figure 2E, F). Interestingly, an abrupt increase in the transfer rate was observed between dT_20_ and dT_30_, suggesting that RPA preferentially binds ssDNA longer than 20 nt under these experimental conditions. Overall, similar to SSB, the ssDNA-bound RPA transferred to competing ssDNA following pseudo-first-order kinetics, and the process displays a nonlinear ssDNA length dependence.

### Real-time SSB and RPA transfer pathway

To elucidate greater details of transfer dynamics, we examined the single-molecule time traces. The pdT_70_-SSB complex by itself showed dynamic *E*_*FRET*_ fluctuation, making it difficult to clearly observe structural dynamics induced by the competing ssDNA (Supplementary Figure S4). Therefore, we used the pdT_40_-SSB complex, which displayed a steady *E*_*FRET*_ state in the absence of competing ssDNA (Supplementary Figure S4). smFRET traces of real-time SSB transfer to the competing strand (dT_40_) showed a sudden one-step drop of *E*_*FRET*_, which also made it challenging to distinguish any intermediate state that might be present (Supplementary Figure S4). Therefore, we employed a different approach to track the intermediate state during strand transfer by using Cy3-labeled competing ssDNA instead of unlabeled ssDNA (Figure 3A).

**Figure 3.**
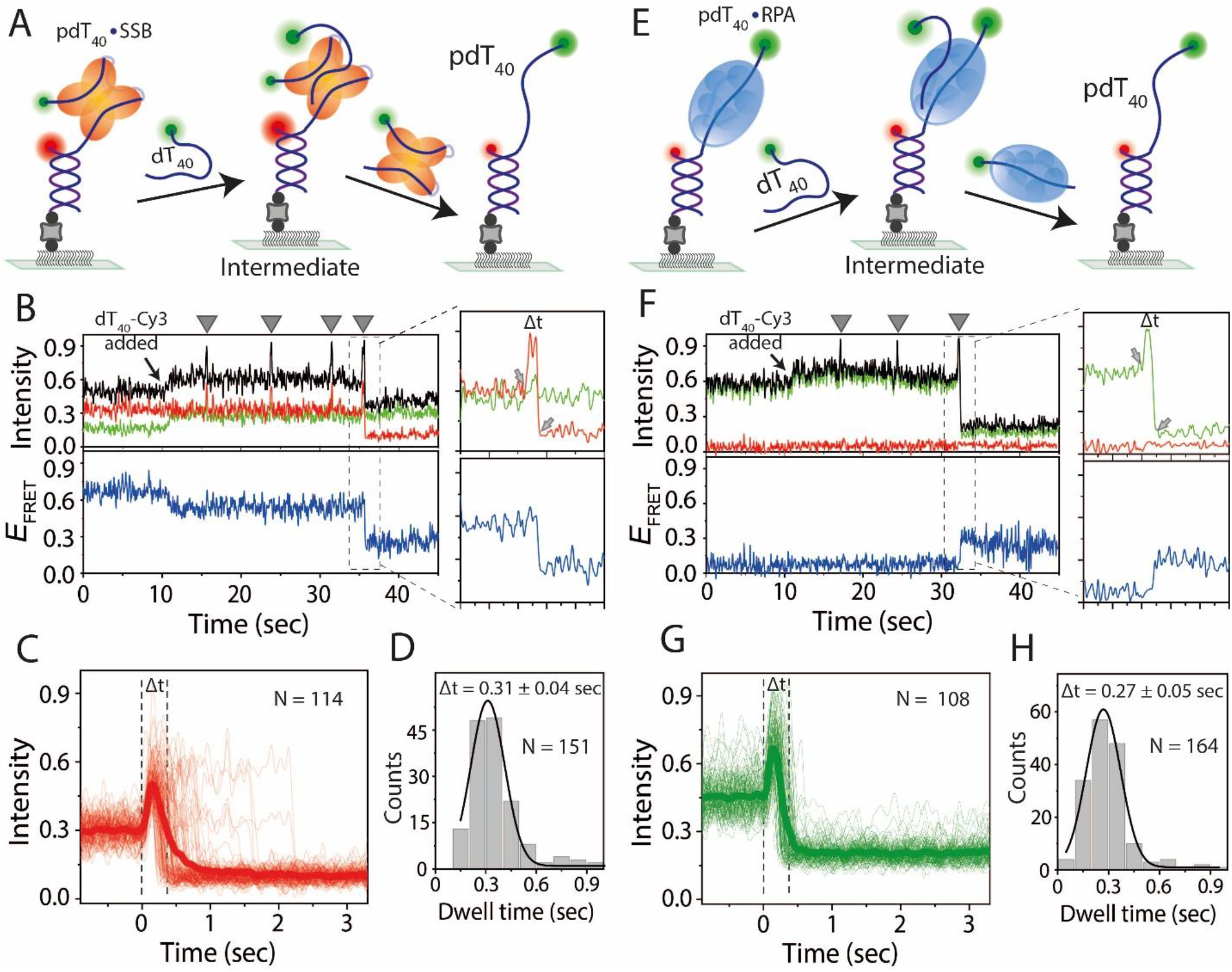
Real-time direct transfer mechanism of SSB and RPA in-between ssDNA. (A, E) Schematics showing SSB (A) or RPA (E) transferring from tethered DNA to Cy3-labeled competing ssDNA. (B, F) Real-time smFRET traces showing SSB (B) and RPA (F) transfer events. Arrows indicate Cy3-ssDNA flow; spikes represent transfer attempts (gray triangles), and boxed regions show successful transfers with intensity and FRET efficiency changes. The background intensity increases after addition of competing Cy3-labeled ssDNA. Black, red, green, and blue lines indicate total intensity, Cy5 intensity, Cy3 intensity, and FRET efficiency, respectively. Two arrows at the zoomed box represent the binding to successful transfer. (C, G) Combined time traces of successful transfers synchronized at the binding event, the moment of intensity increase. The bold line is the average of individuals. (D, H) Gaussian fits of dwell time histograms for successful transfers of SSB (D) and RPA (H), with corresponding dwell times (Δt) indicated (calculated from the zoomed box, time difference between two arrows).

Upon addition of Cy3-labeled dT_40_ (3 nM) to the pdT_40_-SSB complex, the real-time smFRET traces showed an initial background intensity increase (at ~10 sec) due to labeled ssDNA diffusively present in the evanescent field of total internal reflection fluorescence excitation, followed by several Cy3 intensity spikes due to transient association of Cy3-dT_40_ with the surface-tethered pdT_40_-SSB complex until finally, the *E*_*FRET*_ changed to the DNA-alone state, suggesting completion of transfer (Figure 3B). Such spikes reflect the competing ssDNA transiently interacting with the SSB-bound complex, likely forming an intermediate complex for transfer which fails many times prior to achieving a successful transfer. Just before a successful transfer, we observed a sudden intensity increase of Cy5 (due to two Cy3 molecules transferring excitation to a single Cy5). This signature of pre-transfer ternary complex formation is visualized in the combined traces synchronized at the initial moment of Cy5 intensity increase, i.e., at the moment of Cy3-dT_40_ binding (Figure 3C). The average time between binding and complete transfer was ~0.31 sec (Figure 3D).

The strand transfer mechanism of RPA was also examined using Cy3-labeled dT_40_ ssDNA because, like SSB, RPA strand transfer did not show any intermediate *E*_*FRET*_ change when we used unlabeled competing ssDNA (Supplementary Figure S4). Upon addition of Cy3-labeled dT_40_ (3 nM) to the pdT_40_-RPA complex, real-time smFRET traces revealed transient Cy3 intensity spikes (Figure 3E, F), likely representing transient association of competing Cy3-labeled ssDNA with pdT_40_-RPA complex. Successful strand transfer events were characterized by a sudden drop in total intensity, due to the departure of Cy3-dT_40_-RPA complex, and a transition of *E*_*FRET*_ to the DNA-alone state (~0.3). Combined traces synchronized at the binding moment showed intensity spikes followed by intensity drops, confirming the formation of a tripartite intermediate complex during the transfer (Figure 3G). The average time between binding and complete transfer was ~0.27 sec (Figure 3H). Overall, our data showed that both SSB and RPA transfer from one DNA to another occurs through a transient tripartite complex containing both the incumbent DNA strand and the competing DNA strand, and their competition for SSB or RPA can result in multiple failed attempts until a successful transfer.

### Protein exchange dynamics of SSB and RPA on DNA bound complexes

So far, we have used competing ssDNA to study the direct transfer of SSB or RPA from one ssDNA to another. Next, we designed an smFRET assay to quantitatively characterize the exchange of a DNA-bound protein for a protein in solution. We applied labeled proteins to surface-tethered partial-duplex DNA, followed by the addition of unlabeled proteins (Figure 4A, B). It was expected that the initial fluorescence signal that appeared upon labeled protein binding to DNA would disappear over time if unlabeled proteins replaced the labeled.

**Figure 4.**
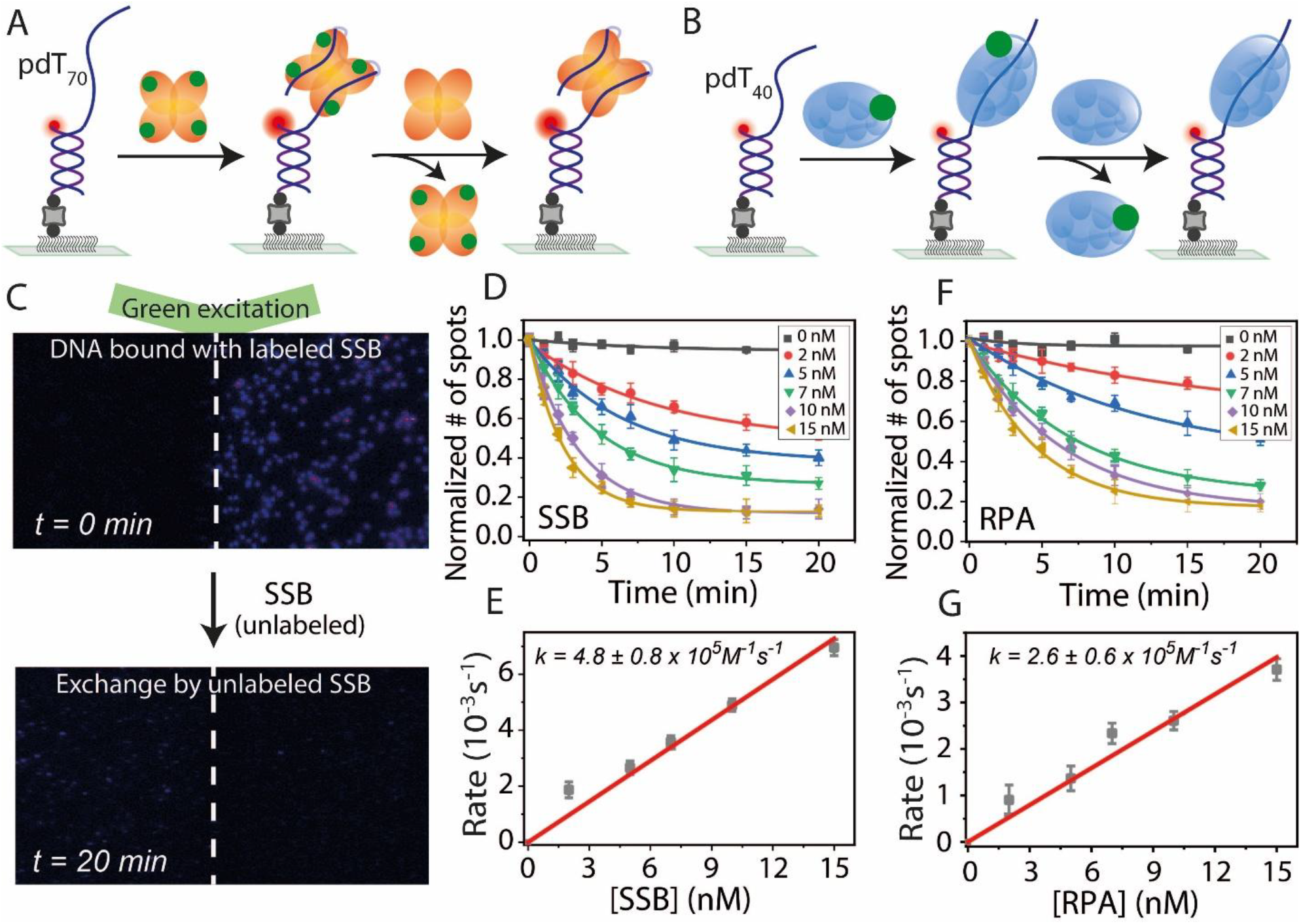
DNA bound protein exchange dynamics of SSB and RPA. (A, B) Schematic representation of SSB (A) and RPA (B) protein-protein exchange. Labeled proteins bound to DNA are replaced by unlabeled proteins. (C) Representative field of view showing labeled SSB at t=0 min (top) and after exchange by unlabeled protein at t=20 min (bottom) under green laser excitation. (D, F) Single-exponential fits of labeled protein disappearance at different unlabeled protein concentrations for SSB (D) and RPA (F). (E, G) Linear fits of protein exchange rates at varying unlabeled protein concentrations for SSB (E) and RPA (G).

After adding Cy3-labeled SSB proteins (2 nM) to the immobilized Cy5-labeled DNA (for initial localization), we observed on average ~300 labeled SSB molecule spots per field of view. A reduction in the number of fluorescent spots was observed over time following the addition of unlabeled SSB proteins (Figure 4C). For kinetic analysis, we quantified the number of fluorescence spots by capturing short movies (~2 s) sequentially at different fields of view to minimize the contribution of photobleaching. To examine the concentration dependence of protein exchange, we titrated unlabeled protein concentrations from 2 nM to 15 nM (Figure 4D). The exchange rates, obtained from single-exponential fits, showed a linear dependence on protein concentration with the bimolecular association rate of 4.8 (± 0.8) 10^5^ M^−1^ s^−1^ for SSB binding to an SSB-DNA complex (Figure 4E). Interestingly, SSB protein exchange rate was lower for shorter ssDNA (40 nt), 3.8 (± 0.3) 10^5^ M^−1^ s^−1^, with a lower exchange fraction (~60%) compared to ~90% for the longer ssDNA of 70 nt at the same protein concentration (Supplementary Figure S5), possibly due to a smaller landing pad size for initial binding of SSB to a ssDNA-SSB complex.

Similar to SSB, we found that surface-tethered DNA bound to MB543-labeled RPA also exhibits protein-protein exchange in a concentration-dependent manner. (Figure 4F), but both the bimolecular exchange rate and the exchange fraction were similar between dT_40_ and dT_70_ ssDNA (2.6 ± 0.6 × 10^5^ M^−1^ s^−1^ for pdT_40_ and 2.7 ± 0.2 × 10^5^ M^−1^ s^−1^ for pdT_70_, ~80% for pdT_40_ and ~75% for pdT_70_) (Figure 4F, G and Supplementary Figure S5). Taken together, these results demonstrate ‘facilitated dissociation’ where dissociation of SSB or RPA from DNA, which is normally very slow (Figure 4D, F), is greatly accelerated by proteins in solution that can replace the incumbents.

### Hetero-protein exchange dynamics between SSB and RPA on ssDNA

Both SSB and RPA are sequence-independent ssDNA-binding proteins, and our study suggests they can undergo homo-protein exchange when bound to DNA (Figure 4). Next, we investigated hetero-protein exchange between SSB and RPA. Although SSB and RPA originate from different organisms, their DNA-binding activities and functions are highly similar (15,49). Observation of hetero-protein exchange would indicate that facilitated dissociation observed in homo-protein exchange does not require specific protein-protein interactions between an incumbent and a competing protein. We added SSB to immobilized DNA, followed by the addition of RPA, or vice versa (Figure 5A). The high-*E*_*FRET*_ peak confirmed SSB binding to ssDNA (pdT_70_). Upon adding RPA, the SSB-bound high-*E*_*FRET*_ peak disappeared and a low-*E*_*FRET*_ peak appeared in its place, confirming that RPA can replace the DNA-bound SSB (Figure 5B). Similarly, SSB replaced DNA-bound RPA, as observed by the transition from low-to high-*E*_*FRET*_.

**Figure 5.**
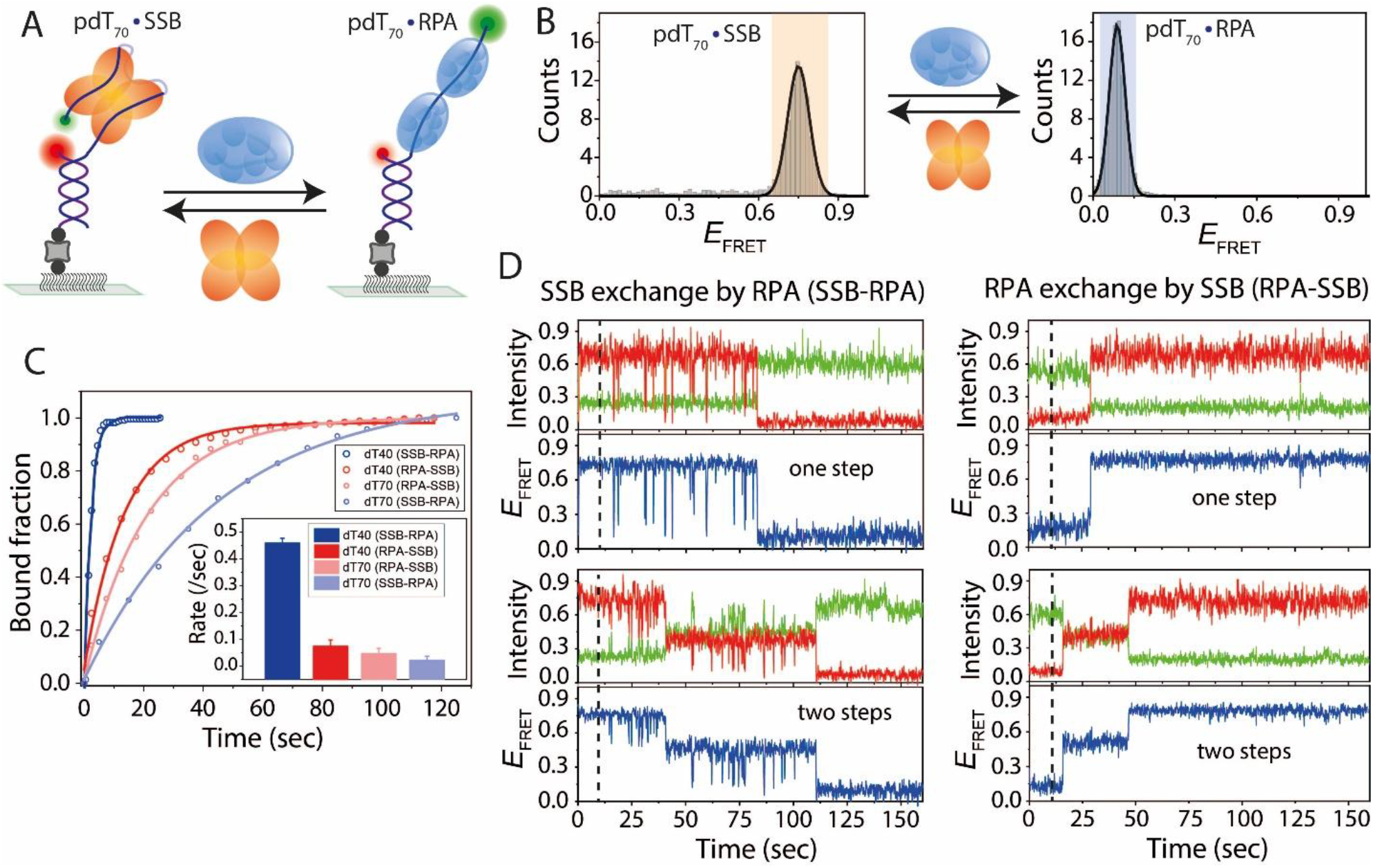
Hetero-protein exchange dynamics between SSB and RPA on ssDNA. (A) Schematic smFRET model of hetero-protein exchange between SSB and RPA. (B) FRET histograms of pdT_70_ showing SSB and RPA binding and their exchange. (C) Single-exponential fits of hetero-protein exchange (SSB to RPA and vice versa) on pdT_40_ and pdT_70_ at 100 nM protein concentrations. (D) Real-time smFRET traces showing SSB-to-RPA and RPA-to-SSB exchanges on pdT_70_, with distinct FRET transition behaviors. Dashed lines indicate the addition of the competing protein.

To study the exchange kinetics, the initial bound fraction was plotted over time (Figure 5C). The overall exchange rate of SSB-to-RPA or RPA-to-SSB was higher on shorter ssDNA (pdT_40_) compared to longer ssDNA (pdT_70_). We further analyzed the kinetics of hetero-protein exchange using smFRET time traces. For the longer ssDNA, two distinct exchange behaviors were observed, while the shorter ssDNA predominantly exhibited one type of behavior (Figure 5D and Supplementary Figure S6). During SSB-to-RPA exchange, SSB-bound high-*E*_*FRET*_ states transitioned through high-to-low *E*_*FRET*_ dynamics as intermediates before stabilizing at an RPA-bound low-*E*_*FRET*_ state, marking completion of exchange. Additionally, on longer ssDNA (dT_70_), a two-step exchange was observed, suggesting that two RPA molecules replace a single SSB molecule in a stepwise manner (48,50). Conversely, during RPA-to-SSB exchange, the RPA-bound low-*E*_*FRET*_ state transitioned in one or two steps to the SSB-bound high-*E*_*FRET*_ state. The two-step transition suggests that two RPA molecules on the ssDNA are sequentially replaced by a single SSB molecule. Altogether, these results demonstrate that SSB and RPA can replace each other on ssDNA even though they must have evolved without an opportunity to interact with each other, indicating that protein exchange does not require specific protein-protein interaction.

### RNA binding and strand transfer dynamics of SSB and RPA

Although SSB and RPA primarily bind to ssDNA, very few studies have reported their ability to bind RNA (40,51), and their biological roles remain poorly understood. Here, we investigated in detail the binding of SSB and RPA to RNA and their associated transfer kinetics. We designed smFRET partial duplex (pd) constructs similar to the DNA constructs, with an overhang consisting of 50 nt of poly-uracil RNA (pdrU_50_) (Figure 6A). Upon addition of SSB (10 nM) to pdrU_50_ in a buffer containing 100 mM NaCl, the low-*E*_*FRET*_ peak (~0.2) corresponding to RNA-only shifted to a mid-*E*_*FRET*_ peak (~0.6), suggesting that SSB binding to RNA can also induce wrapping of RNA on the SSB surface (Figure 6B). Similar to the experiments with DNA, we introduced competing (50-nt of poly-uracil, rU_50_) RNA into the sample chamber to study SSB strand transfer from bound RNA to RNA from solution. Over time, the mid-*E*_*FRET*_ bound fraction corresponding to SSB-bound RNA decreased, while the low-*E*_*FRET*_ RNA-only population increased. The bound fraction plotted over time at various rU_50_ RNA concentrations (1 nM to 20 nM) fitted well with single-exponential decay curves, consistent with pseudo-first-order kinetics as observed with DNA (Figure 6C). The transfer rates exhibited a linear dependence on rU_50_ concentration, yielding a bimolecular rate constant of 2.1 (± 0.2) × 10^5^ M^−1^ s^−1^ (Figure 6D).

**Figure 6.**
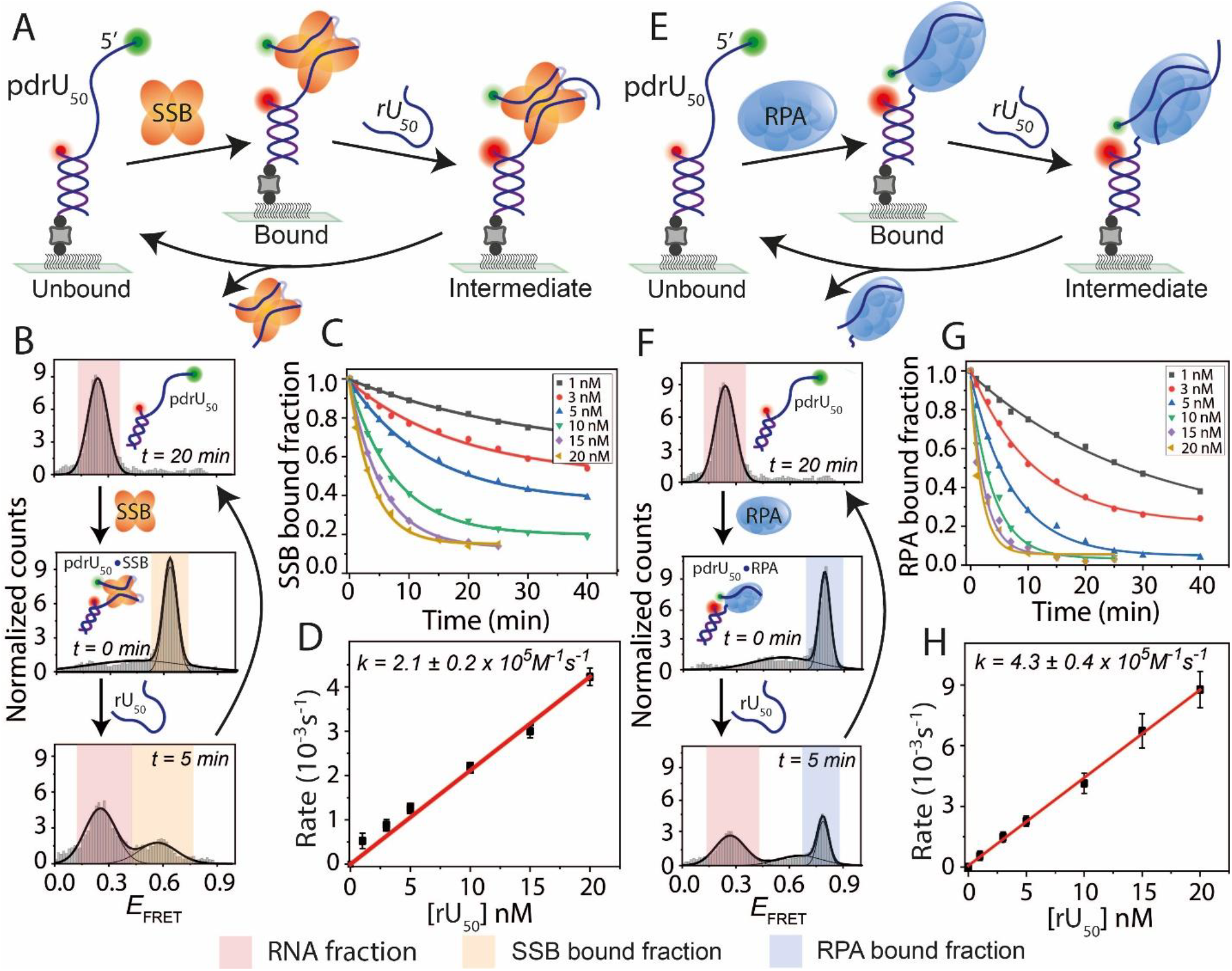
RNA binding and strand transfer dynamics of SSB and RPA. (A, E) Schematics of smFRET constructs with a partial duplex containing a 50-nt poly-uracil overhang (pdrU_50_). SSB (A) or RPA (E) binds pdrU_50_, followed by transfer to competing RNA (rU_50_). (B, F) FRET histograms of pdrU_50_ before (top, light pink) and after (middle, light orange) SSB (B) or RPA (F) binding, and during SSB or RPA transfer (bottom) upon rU_50_ addition at indicated times. Time, t=20 min (top histogram) represents the complete strand transfer of bound SSB or RPA. (C, G) Single-exponential fits of protein-bound fractions during RNA-induced transfer at different rU_50_ concentrations for SSB (C) and RPA (G). (D, H) Linear fits of transfer rates at varying rU_50_ concentrations for SSB (D) and RPA (H).

The same RNA binding and direct strand transfer assays were performed for RPA (Figure 6E). Interestingly, upon addition of RPA (10 nM) to pdrU_50_ in a buffer containing 100 mM NaCl, unlike DNA, the RPA-bound RNA exhibited a high-*E*_*FRET*_ state (~0.8), indicating that RNA wraps around the RPA protein (Figure 6F). Similar to SSB, RPA-bound RNA followed pseudo-first-order kinetics, with transfer rates displaying a linear dependence on rU_50_ concentration with a bimolecular rate constant of 4.3 (± 0.4) × 10^5^ M^−1^ s^−1^ (Figure 6G, H). Altogether, these results demonstrate that both SSB and RPA bind RNA at nanomolar concentrations and efficiently transfer to competing RNA via pseudo-first-order kinetics, mirroring their behavior with ssDNA.

### Binding competition between DNA and RNA for SSB or RPA

Our results showed that both SSB and RPA can bind to DNA and RNA. Next, we investigated the binding competition between DNA and RNA for these proteins. To do this, we immobilized both DNA (pdT_70_) and RNA (pdrU_50_) on the same surface, ensuring nearly equal populations (Figure 7A). The *E*_*FRET*_ peaks for DNA and RNA alone were both located in the low-*E*_*FRET*_ region (~0.2–0.3) (Figure 7B). However, the SSB-bound *EFRET* peaks were clearly distinguishable, with mid-*EFRET* representing RNA-bound SSB and high-*E*_*FRET*_ representing DNA-bound SSB. Interestingly, upon adding a competing RNA strand (rU_50_), the SSB-bound RNA population at mid-*E*_*FRET*_ completely shifted to the low-*E*_*FRET*_ RNA-only state, while the DNA-bound SSB population at high-*E*_*FRET*_ remained unchanged. Even with the addition of a high concentration of RNA (10 µM), the SSB-bound DNA high-*E*_*FRET*_ population was unaffected. Conversely, the addition of ssDNA (dT_60_) caused both mid- and high-*E*_*FRET*_ peaks to shift entirely to the low-*E*_*FRET*_ DNA- and RNA-only state, indicating complete strand transfer of the bound protein. Therefore, although SSB can also bind RNA, DNA outcompetes RNA in its ability to take away SSB from DNA or RNA.

**Figure 7.**
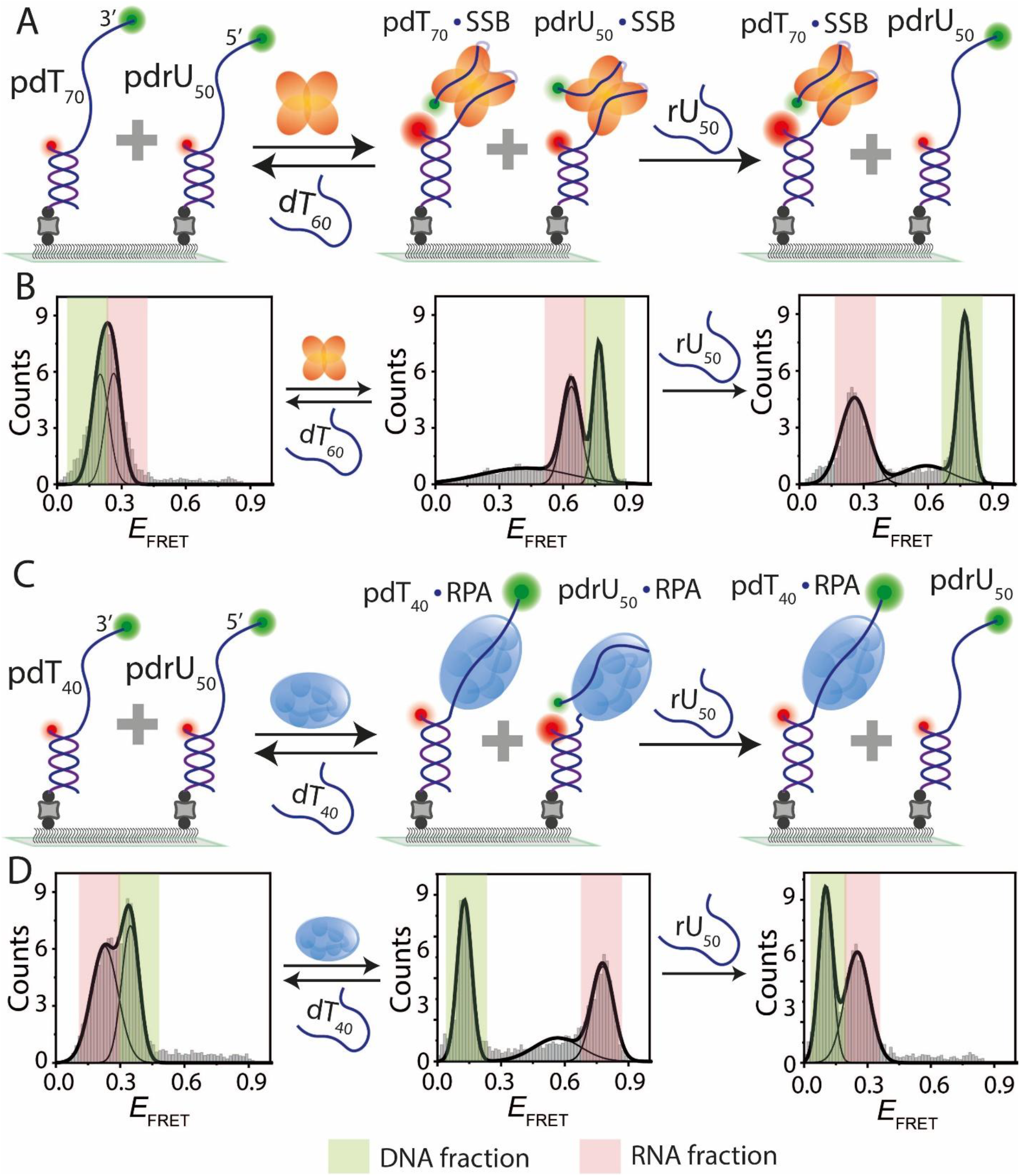
Binding competition between DNA and RNA for SSB and RPA. (A, C) Schematic of smFRET constructs with co-immobilized DNA (pdT_70_/pdT_40_) and RNA (pdrU_50_). SSB (A) or RPA (C) binds to both DNA and RNA, with subsequent transfer to competing ssDNA (dT_60_ for SSB, dT_40_ for RPA) or rU_50_. (B, D) FRET histograms showing DNA and RNA before and after SSB (B) or RPA (D) binding. Competing ssDNA facilitates complete transfer of protein from RNA and DNA, whereas competing RNA transfers only RNA-bound protein. Colors represent respective bound fractions.

A similar competition assay was performed with RPA (Figure 7C). RPA bound to DNA and RNA produced well-separated low- and high-*E*_*FRET*_ peaks (Figure 7D). The RNA-bound RPA high-*E*_*FRET*_ population completely shifted to the low-*E*_*FRET*_ RNA-only state upon the addition of RNA (rU_50_), whereas the DNA-bound *E*_*FRET*_ population remained unaffected even after adding 10 µM RNA. Upon adding ssDNA (dT_60_), both *E*_*FRET*_ peaks shifted completely to the original DNA- and RNA-only state, similar to SSB. In summary, the binding competition assays for SSB and RPA show that, although both proteins can bind RNA, they strongly prefer binding to DNA over RNA.

## DISCUSSION

Using single-molecule FRET assays, we uncovered fine details on how these proteins orchestrate nucleic acid transactions with remarkable adaptability. Our findings illuminate the competitive binding, direct transfer, and exchange processes that enable SSB and RPA to sustain genome stability and contend with RNA metabolism in fluctuating cellular environments.

### Mechanistic insights into strand transfer and protein exchange dynamics

Strand transfer is a fundamental process that ensures the efficiency and fidelity of nucleic acid metabolism, particularly during replication, where ssDNA-binding proteins must rapidly adapt to changing substrates and redistribute to maintain polymerase activity (7,15). This process enables proteins to efficiently replace pre-bound complexes or move between ssDNA regions without dissociating completely into free protein intermediates. Here, we provide direct visualization of the strand transfer mechanism, building on prior research that extensively characterized the ssDNA-binding behavior of SSB and RPA proteins (Figure 3). Our findings reveal that strand transfer in both proteins proceed through a ternary intermediate complex, where the protein simultaneously interacts with at least two ssDNA molecules. This direct transfer mechanism bypasses the need for a protein-free intermediate, minimizing the risk of exposing ssDNA to nucleases or other damaging agents. Repeated transfer attempts, observed as transient smFRET intensity spikes, indicate the formation of unstable intermediate complexes that continue until incoming strand wins the battle. This strand transfer mechanism likely involves partial opening of the protein-DNA complex or a flexible binding mode that accommodates additional ssDNA. Additionally, the diffusion properties of SSB and RPA enable them to migrate along ssDNA (22,26,27), facilitating engagement with competing strands. The process is highly concentration-dependent, with strand transfer rates increasing linearly with competing ssDNA concentration, indicating that a single competing strand is involved in the rate-determining step, and the overall reaction rate is governed by diffusion-controlled encounters.

Strand transfer efficiency is notably influenced by competing ssDNA’s length, with longer strands (≥60 nt for SSB and ≥40 nt for RPA) significantly enhancing transfer efficiency (Figure 1, 2). Longer ssDNA segments are likely to promote stronger protein engagement, leading to greater destabilization of the pre-bound complex and facilitating more efficient transfer. These results suggest that replication forks, which naturally generate longer ssDNA regions, provide an optimal environment for strand transfer *in vivo*. While our use of poly-thymine sequences removed sequence specificity, it is important to note that the efficiency of direct transfer may vary depending on DNA sequences composition. Notably, sequence-specific proteins such as POT1, which share similar binding affinities with SSB and RPA, do not undergo strand transfer (52). This observation underscores the versatility of SSB and RPA in adapting to different DNA substrates. Understanding these mechanisms sheds light on how strand transfer supports the rapid redistribution and replacement of ssDNA-binding proteins, ensuring genome integrity during critical cellular processes.

### Protein exchange dynamics of SSB and RPA on ssDNA

SSB and RPA are essential for maintaining nucleic acid integrity by enabling timely protein exchange during DNA metabolic processes. Both proteins bind tightly to ssDNA to protect it, but they must also be displaced efficiently to allow access for other DNA-processing factors. This rapid protein exchange is driven by their diffusion properties, where bound proteins are replaced by the binding of a protein in solution to a ssDNA segment transiently exposed by the diffusional migration of an incumbent protein. Previous studies have proposed that the concentration-dependent turnover of dsDNA-binding proteins facilitates dissociation and rapid rebinding of proteins on DNA (53). Additionally, DNA curtain assays have demonstrated that eGFP-tagged RPA rapidly exchanges with untagged RPA in a concentration-dependent manner (32,43). Consistent with these findings, our data show that both SSB and RPA undergo rapid exchange with freely diffusing proteins in a concentration-dependent manner (Figure 4). Even when bound to short ssDNA substrates, where only a single SSB tetramer or one to two RPA monomers could bind, rapid protein exchange was still observed, reflecting intrinsic exchange properties of both proteins. This behavior supports the idea that SSB and RPA can undergo rapid protein exchange, particularly given the high intracellular concentrations of DNA as well as SSB (~0.05–0.3 µM) and RPA (~2 µM), which are higher than the concentrations used in our experiments (54,55).

In addition to homo-protein exchange, SSB and RPA also exhibit hetero-protein exchange, replacing each other on ssDNA in a bidirectional manner (Figure 5). This exchange demonstrates that SSB and RPA can replace each other on ssDNA *in vitro*, highlighting similarities in their DNA binding dynamics despite originating from different organisms. Trimeric RPA was observed to replace SSB tetramers or vice versa, with dynamics modulated by ssDNA length, reflecting differences in their binding strategies. Our single-molecule time traces provided direct evidence of hetero-protein exchange between SSB and RPA, revealing distinct exchange kinetics. RPA-induced SSB exchange was observed as fluctuations in FRET intensity, while reverse exchange dynamics were not observed, likely due to differences in the modes of protein binding. The exchange kinetics were dependent on ssDNA length, with longer ssDNA showing a one- or two-step exchange process. This offers insight into the complex interplay between these proteins, especially when binding sites overlap or higher-order nucleoprotein structures form.

These findings have important implications for genome maintenance, as SSB and RPA must rapidly adapt to changing DNA structures and respond to challenges such as DNA damage and replication stress (11,14,33,56). Protein exchange plays a central role in regulating DNA and RNA dynamics, ensuring efficient turnover of bound factors and facilitating the recruitment of DNA-processing proteins. (14,15). For instance, in telomere regulation, RPA is replaced by POT1 to shift from replication to telomere protection (57) while in homologous recombination, RAD51 replaces RPA to promote strand invasion and repair (32,58). Additionally, RNA-binding proteins undergo dynamic exchange to regulate RNA stability and processing during transcription and to modulate DNA damage signaling by displacing inappropriate proteins (59,60). These observations highlight the adaptability of ssDNA-binding proteins in maintaining nucleic acid homeostasis. The ability of SSB and RPA to undergo both homo- and hetero-protein exchange suggests that similar exchange-based regulatory mechanisms could be broadly relevant across different nucleic acid-processing pathways.

### RNA interactions and functional versality of SSB and RPA

In addition to their established roles in ssDNA metabolism, our study reveals that SSB and RPA also interact with RNA, undergo strand transfer to competing RNA molecules (Figure 6). However, competitive binding assays demonstrated a strong preference for DNA over RNA (Figure 7), reinforcing their primary functions in DNA transactions. This dual functionality highlights their ability to engage with both DNA and RNA substrates, particularly in cellular contexts where these nucleic acids coexist. Our findings suggest that RNA and ssDNA compete for binding to SSB and RPA, potentially influencing key cellular processes such as transcription-coupled repair, RNA processing, and translation. RNA molecules may displace DNA-bound proteins, or vice versa, thereby modulating cellular responses to genomic instability. The physical and functional interaction of SSB with RNA polymerase further supports its involvement in co-transcriptional RNA processes (36,37). While SSB may transiently interact with nascent RNA, its stronger preference for DNA may limit RNA binding under many conditions. In contrast, RPA is closely associated with R-loops *in vivo* and exhibits a stronger affinity for co-transcriptional RNA, potentially promoting the formation of DNA-RNA hybrids or R-loops, which play critical roles in gene expression regulation and genome stability (38–40).

Recent studies have shown that multiple RNA- and DNA-binding proteins can directly transfer between polynucleotides without requiring a free protein intermediate (30). This direct transfer capability, also observed in our study, likely represents a general property of nucleic acid-binding proteins. Such mechanisms are essential for RNA-binding proteins to recruit factors to chromatin and for DNA-binding proteins to efficiently locate their targets on nucleosomal DNA. Furthermore, direct transfer from nascent RNA to DNA may explain why many DNA-binding proteins also exhibit RNA-binding capabilities, reducing the complexity of their search for binding sites and enabling dynamic regulation of nucleic acid interactions. These findings expand the functional repertoire of SSB and RPA, demonstrating their adaptability in managing both RNA and DNA substrates. Their dual roles in genome and transcriptome maintenance underscore their importance in coordinating cellular processes that require DNA and RNA interplay, especially during transcriptional stress or genomic instability.

### Broader implications and future directions

The mechanistic insights presented here enhance our understanding of the evolutionary adaptations of SSB and RPA to diverse cellular environments. Their ability to mediate strand transfer, facilitate rapid protein exchange, and interact with RNA emphasizes their essential roles in genome stability and RNA metabolism. The observed preference for DNA over RNA and the context-dependent dynamics of protein exchange suggest a finely tuned regulatory system that responds to cellular needs. Future studies should focus on unraveling the structural basis of these interactions, particularly the conformational changes during strand transfer and protein exchange. Investigating these mechanisms under physiological conditions, such as during transcription-coupled repair or replication stress responses, will further clarify their biological relevance. Exploring sequence-specific strand transfer and RNA interactions could reveal additional layers of functional specificity, particularly in the context of telomere maintenance, homologous recombination, and RNA-mediated chromatin regulation. By integrating these insights, we advance the broader understanding of protein-nucleic acid interactions and pave the way for targeted therapeutic strategies to address conditions involving genome instability and dysregulated RNA metabolism.

## ACKNOWLEDGEMENTS

We thank all the members of the Sua Myong and Taekjip Ha laboratories for their helpful comments and constructive criticisms. This article is subject to HHMI’s Open Access to Publications policy. HHMI lab heads have previously granted a nonexclusive CC BY 4.0 license to the public and a sublicensable license to HHMI in their research articles. Pursuant to those licenses, the author-accepted manuscript of this article can be made freely available under a CC BY 4.0 license immediately upon publication.

## Author contributions

Tapas Paul: Conceptualization, Data curation, Data analysis, Investigation, Methodology, Project administration, Validation, Visualization, Writing-original draft, Writing-review and editing. I-Ren Lee: Conceptualization, Data curation, Formal analysis, Methodology. Sushil Pangeni: Conceptualization, Discussion. Fahad Rashid: SSB protein Purification and labeling. Olivia Yang: Conceptualization. Edwin Antony: RPA protein purification and labeling, Funding acquisition. James M. Berger: Supervision. Sua Myong: Funding acquisition, Supervision. Taekjip Ha: Conceptualization, Supervision, Funding acquisition, Writing-review and editing.

## REFERENCES

1. Dueva, R. and Iliakis, G. (2020) Replication protein A: a multifunctional protein with roles in DNA replication, repair and beyond. NAR cancer, 2, zcaa022.

2. Broderick, S., Rehmet, K., Concannon, C. and Nasheuer, H.-P. (2010) Eukaryotic single-stranded DNA binding proteins: central factors in genome stability. Genome stability and human diseases, 143–163.

3. Antony, E. and Lohman, T.M. (2019), Seminars in cell & developmental biology. Elsevier, Vol. 86, pp. 102–111.

4. Meyer, R.R. and Laine, P.S. (1990) The single-stranded DNA-binding protein of Escherichia coli. Microbiological reviews, 54, 342–380.

5. Wang, Y.-R., Guo, T.-T., Zheng, Y.-T., Lai, C.-W., Sun, B., Xi, X.-G. and Hou, X.-M. (2021) Replication protein A plays multifaceted roles complementary to specialized helicases in processing G-quadruplex DNA. Iscience, 24.

6. Brill, S.J. and Bastin-Shanower, S. (1998) Identification and characterization of the fourth single-stranded-DNA binding domain of replication protein A. Molecular and cellular biology, 18, 7225–7234.

7. Lohman, T.M. and Ferrari, M.E. (1994) Escherichia coli single-stranded DNA-binding protein: multiple DNA-binding modes and cooperativities. Annual review of biochemistry, 63, 527–570.

8. Morten, M.J., Peregrina, J.R., Figueira-Gonzalez, M., Ackermann, K., Bode, B.E., White, M.F. and Penedo, J.C. (2015) Binding dynamics of a monomeric SSB protein to DNA: a single-molecule multi-process approach. Nucleic acids research, 43, 10907–10924.

9. Pangeni, S., Biswas, G., Kaushik, V., Kuppa, S., Yang, O., Lin, C.-T., Mishra, G., Levy, Y., Antony, E. and Ha, T. (2024) Rapid long-distance migration of RPA on single stranded DNA occurs through intersegmental transfer utilizing multivalent interactions. Journal of Molecular Biology, 436, 168491.

10. Bochkareva, E., Belegu, V., Korolev, S. and Bochkarev, A. (2001) Structure of the major single-stranded DNA-binding domain of replication protein A suggests a dynamic mechanism for DNA binding. The EMBO journal.

11. Thrall, E.S., Piatt, S.C., Chang, S. and Loparo, J.J. (2022) Replication stalling activates SSB for recruitment of DNA damage tolerance factors. Proceedings of the National Academy of Sciences, 119, e2208875119.

12. Spenkelink, L.M., Lewis, J.S., Jergic, S., Xu, Z.-Q., Robinson, A., Dixon, N.E. and van Oijen, A.M. (2019) Recycling of single-stranded DNA-binding protein by the bacterial replisome. Nucleic acids research, 47, 4111–4123.

13. Pike, A.M., Friend, C.M. and Bell, S.P. (2023) Distinct RPA functions promote eukaryotic DNA replication initiation and elongation. Nucleic Acids Research, 51, 10506–10518.

14. Maréchal, A. and Zou, L. (2015) RPA-coated single-stranded DNA as a platform for post-translational modifications in the DNA damage response. Cell research, 25, 9–23.

15. Shereda, R.D., Kozlov, A.G., Lohman, T.M., Cox, M.M. and Keck, J.L. (2008) SSB as an organizer/mobilizer of genome maintenance complexes. Critical reviews in biochemistry and molecular biology, 43, 289–318.

16. Acharya, A., Kasaciunaite, K., Göse, M., Kissling, V., Guérois, R., Seidel, R. and Cejka, P. (2021) Distinct RPA domains promote recruitment and the helicase-nuclease activities of Dna2. Nature Communications, 12, 6521.

17. Ha, T., Kozlov, A.G. and Lohman, T.M. (2012) Single-molecule views of protein movement on single-stranded DNA. Annual review of biophysics, 41, 295–319.

18. Plaza-GA, I., Lemishko, K.M., Crespo, R., Truong, T.Q., Kaguni, L.S., Cao-García, F.J., Ciesielski, G.L. and Ibarra, B. (2023) Mechanism of strand displacement DNA synthesis by the coordinated activities of human mitochondrial DNA polymerase and SSB. Nucleic acids research, 51, 1750–1765.

19. Cerron, F., de Lorenzo, S., Lemishko, K.M., Ciesielski, G.L., Kaguni, L.S., Cao, F.J. and Ibarra, B. (2019) Replicative DNA polymerases promote active displacement of SSB proteins during lagging strand synthesis. Nucleic acids research, 47, 5723–5734.

20. Sansam, C.L. and Pezza, R.J. (2015) Connecting by breaking and repairing: mechanisms of DNA strand exchange in meiotic recombination. The FEBS journal, 282, 2444–2457.

21. Qi, Z. and Greene, E.C. (2016) Visualizing recombination intermediates with single-stranded DNA curtains. Methods, 105, 62–74.

22. Roy, R., Kozlov, A.G., Lohman, T.M. and Ha, T. (2009) SSB protein diffusion on single-stranded DNA stimulates RecA filament formation. Nature, 461, 1092–1097.

23. Lee, K.S., Marciel, A.B., Kozlov, A.G., Schroeder, C.M., Lohman, T.M. and Ha, T. (2014) Ultrafast redistribution of E. coli SSB along long single-stranded DNA via intersegment transfer. Journal of molecular biology, 426, 2413–2421.

24. Mishra, G., Bigman, L.S. and Levy, Y. (2020) ssDNA diffuses along replication protein A via a reptation mechanism. Nucleic acids research, 48, 1701–1714.

25. Zhou, R., Kozlov, A.G., Roy, R., Zhang, J., Korolev, S., Lohman, T.M. and Ha, T. (2011) SSB functions as a sliding platform that migrates on DNA via reptation. Cell, 146, 222–232.

26. Nguyen, B., Sokoloski, J., Galletto, R., Elson, E.L., Wold, M.S. and Lohman, T.M. (2014) Diffusion of human replication protein A along single-stranded DNA. Journal of molecular biology, 426, 3246–3261.

27. Gorman, J. and Greene, E.C. (2008) Visualizing one-dimensional diffusion of proteins along DNA. Nature structural & molecular biology, 15, 768–774.

28. Lohman, T.M. and Overman, L.B. (1985) Two binding modes in Escherichia coli single strand binding protein-single stranded DNA complexes. Modulation by NaCl concentration. Journal of Biological Chemistry, 260, 3594–3603.

29. Kozlov, A.G. and Lohman, T.M. (2002) Kinetic mechanism of direct transfer of Escherichia coli SSB tetramers between single-stranded DNA molecules. Biochemistry, 41, 11611–11627.

30. Hemphill, W.O., Voong, C.K., Fenske, R., Goodrich, J.A. and Cech, T.R. (2023) Multiple RNA-and DNA-binding proteins exhibit direct transfer of polynucleotides with implications for target-site search. Proceedings of the National Academy of Sciences, 120, e2220537120.

31. Menetski, J.P. and Kowalczykowski, S.C. (1987) Transfer of recA protein from one polynucleotide to another. Kinetic evidence for a ternary intermediate during the transfer reaction. Journal of Biological Chemistry, 262, 2085–2092.

32. Gibb, B., Ling, F.Y., Gergoudis, S.C., Kwon, Y., Niu, H., Sung, P. and Greene, E.C. (2014) Concentration-dependent exchange of replication protein A on single-stranded DNA revealed by single-molecule imaging. PLoS ONE, 9, e87922.

33. Zou, Y., Liu, Y., Wu, X. and Shell, S.M. (2006) Functions of human replication protein A (RPA): from DNA replication to DNA damage and stress responses. Journal of cellular physiology, 208, 267–273.

34. Morten, M.J., Gamsjaeger, R., Cubeddu, L., Kariawasam, R., Peregrina, J., Penedo, J.C. and White, M.F. (2017) High-affinity RNA binding by a hyperthermophilic single-stranded DNA-binding protein. Extremophiles, 21, 369–379.

35. Xiao, Y., Jiang, Z., Zhang, M., Zhang, X., Gan, Q., Yang, Y., Wu, P., Feng, X., Ni, J. and Dong, X. (2023) The canonical single-stranded DNA-binding protein is not an essential replication factor but an RNA chaperon in Saccharolobus islandicus. Iscience, 26.

36. Davydova, E.K. and Rothman-Denes, L.B. (2003) Escherichia coli single-stranded DNA-binding protein mediates template recycling during transcription by bacteriophage N4 virion RNA polymerase. Proceedings of the National Academy of Sciences, 100, 9250–9255.

37. Richard, D.J., Bell, S.D. and White, M.F. (2004) Physical and functional interaction of the archaeal single-stranded DNA-binding protein SSB with RNA polymerase. Nucleic acids research, 32, 1065–1074.

38. Mazina, O.M., Somarowthu, S., Kadyrova, L.Y., Baranovskiy, A.G., Tahirov, T.H., Kadyrov, F.A. and Mazin, A.V. (2020) Replication protein A binds RNA and promotes R-loop formation. Journal of Biological Chemistry, 295, 14203–14213.

39. Yasuhara, T., Kato, R., Hagiwara, Y., Shiotani, B., Yamauchi, M., Nakada, S., Shibata, A. and Miyagawa, K. (2018) Human Rad52 promotes XPG-mediated R-loop processing to initiate transcription-associated homologous recombination repair. Cell, 175, 558–570.e511.

40. Nguyen, H.D., Yadav, T., Giri, S., Saez, B., Graubert, T.A. and Zou, L. (2017) Functions of replication protein A as a sensor of R loops and a regulator of RNaseH1. Molecular cell, 65, 832–847. e834.

41. Paul, T., Voter, A.F., Cueny, R.R., Gavrilov, M., Ha, T., Keck, J.L. and Myong, S. (2020) E. coli Rep helicase and RecA recombinase unwind G4 DNA and are important for resistance to G4-stabilizing ligands. Nucleic acids research, 48, 6640–6653.

42. Paul, T., Yang, L., Lee, C.-Y. and Myong, S. (2024) Simultaneous probing of transcription, G-quadruplex, and R-loop. Methods in Enzymology, 705, 377–396.

43. Pokhrel, N., Origanti, S., Davenport, E.P., Gandhi, D., Kaniecki, K., Mehl, R.A., Greene, E.C., Dockendorff, C. and Antony, E. (2017) Monitoring Replication Protein A (RPA) dynamics in homologous recombination through site-specific incorporation of non-canonical amino acids. Nucleic acids research, 45, 9413–9426.

44. Lee, H.-T., Sanford, S., Paul, T., Choe, J., Bose, A., Opresko, P.L. and Myong, S. (2020) Position-dependent effect of guanine base damage and mutations on telomeric G-quadruplex and telomerase extension. Biochemistry, 59, 2627–2639.

45. Paul, T. and Myong, S. (2022) Protocol for generation and regeneration of PEG-passivated slides for single-molecule measurements. STAR protocols, 3, 101152.

46. Paul, T., Ha, T. and Myong, S. (2021) Regeneration of PEG slide for multiple rounds of single-molecule measurements. Biophysical journal, 120, 1788–1799.

47. Lohman, T.M., Overman, L.B. and Datta, S. (1986) Salt-dependent changes in the DNA binding co-operativity of Escherichia coli single strand binding protein. Journal of molecular biology, 187, 603–615.

48. Chadda, R., Kaushik, V., Ahmad, I.M., Deveryshetty, J., Holehouse, A.S., Sigurdsson, S.T., Biswas, G., Levy, Y., Bothner, B. and Cooley, R.B. (2024) Partial wrapping of single-stranded DNA by replication protein A and modulation through phosphorylation. Nucleic acids research, 52, 11626–11640.

49. Wold, M.S. (1997) Replication protein A: a heterotrimeric, single-stranded DNA-binding protein required for eukaryotic DNA metabolism. Annual review of biochemistry, 66, 61–92.

50. Yates, L.A., Aramayo, R.J., Pokhrel, N., Caldwell, C.C., Kaplan, J.A., Perera, R.L., Spies, M., Antony, E. and Zhang, X. (2018) A structural and dynamic model for the assembly of Replication Protein A on single-stranded DNA. Nature communications, 9, 5447.

51. Wang, Q.-M., Yang, Y.-T., Wang, Y.-R., Gao, B., Xi, X. and Hou, X.-M. (2019) Human replication protein A induces dynamic changes in single-stranded DNA and RNA structures. Journal of Biological Chemistry, 294, 13915–13927.

52. Paul, T., Liou, W., Cai, X., Opresko, P.L. and Myong, S. (2021) TRF2 promotes dynamic and stepwise looping of POT1 bound telomeric overhang. Nucleic Acids Research, 49, 12377–12393.

53. Graham, J.S., Johnson, R.C. and Marko, J.F. (2011) Concentration-dependent exchange accelerates turnover of proteins bound to double-stranded DNA. Nucleic acids research, 39, 2249–2259.

54. Ghaemmaghami, S., Huh, W.-K., Bower, K., Howson, R.W., Belle, A., Dephoure, N., O’Shea, E.K. and Weissman, J.S. (2003) Global analysis of protein expression in yeast. Nature, 425, 737–741.

55. Williams, K., Murphy, J. and Chase, J. (1984) Characterization of the structural and functional defect in the Escherichia coli single-stranded DNA binding protein encoded by the ssb-1 mutant gene. Expression of the ssb-1 gene under lambda pL regulation. Journal of Biological Chemistry, 259, 11804–11811.

56. Morse, M., Roby, F.N., Kinare, M., McIsaac, J., Williams, M.C. and Beuning, P.J. (2023) DNA damage alters binding conformations of E. coli single-stranded DNA-binding protein. Biophysical Journal, 122, 3950–3958.

57. Ray, S., Bandaria, J.N., Qureshi, M.H., Yildiz, A. and Balci, H. (2014) G-quadruplex formation in telomeres enhances POT1/TPP1 protection against RPA binding. Proceedings of the National Academy of Sciences, 111, 2990–2995.

58. Li, X. and Heyer, W.-D. (2008) Homologous recombination in DNA repair and DNA damage tolerance. Cell research, 18, 99–113.

59. Hentze, M.W., Castello, A., Schwarzl, T. and Preiss, T. (2018) A brave new world of RNA-binding proteins. Nature reviews Molecular cell biology, 19, 327–341.

60. Shen, H. and Green, M.R. (2004) A pathway of sequential arginine-serine-rich domain-splicing signal interactions during mammalian spliceosome assembly. Molecular cell, 16, 363–373.

