## Supplementary Information for "Mechanistic insights into direct DNA and RNA strand transfer and dynamic protein exchange of SSB and RPA"

Supplementary information includes:
Supplementary Table 1
Supplementary Figures, S1-S6

**Supplementary Table 1.** DNA Oligonucleotides (5’ to 3’) used in this experiment.

| | |
|---|---|
| 18mer-poly-thymine_Cy3 | TGG CGA CGG CAG CGA GGC $(T)_n$-/Cy3/ (n = 40, 70) |
| Cy5-18mer-Bio | /Cy5/-GCC TCG CTG CCG TCG CCA-/Bio/ |
| Unlabeled Poly-thymine | $(T)_n$ (n = 40, 60) |
| Poly-thymine_Cy3 | $(T)_{40}$-Cy3 |
| 18mer-poly-thymine | TGG CGA CGG CAG CGA GGC $(T)_n$ (n = 40, 70) |
| 18mer-poly-uracil _Amine | /Amine/-$(rU)_{50}$ rGrCrC rUrCrG rCrUrG rCrCrG rUrCrG rCrCrA |
| Bio -18mer-Cy5 | /Bio/-rUrGrG rCrGrA rCrGrG rCrArG rCrGrA rGrGrC-/Cy5/ |
| Unlabeled Poly-uracil | $(U)_{50}$ |

**Supplementary Figure 1**

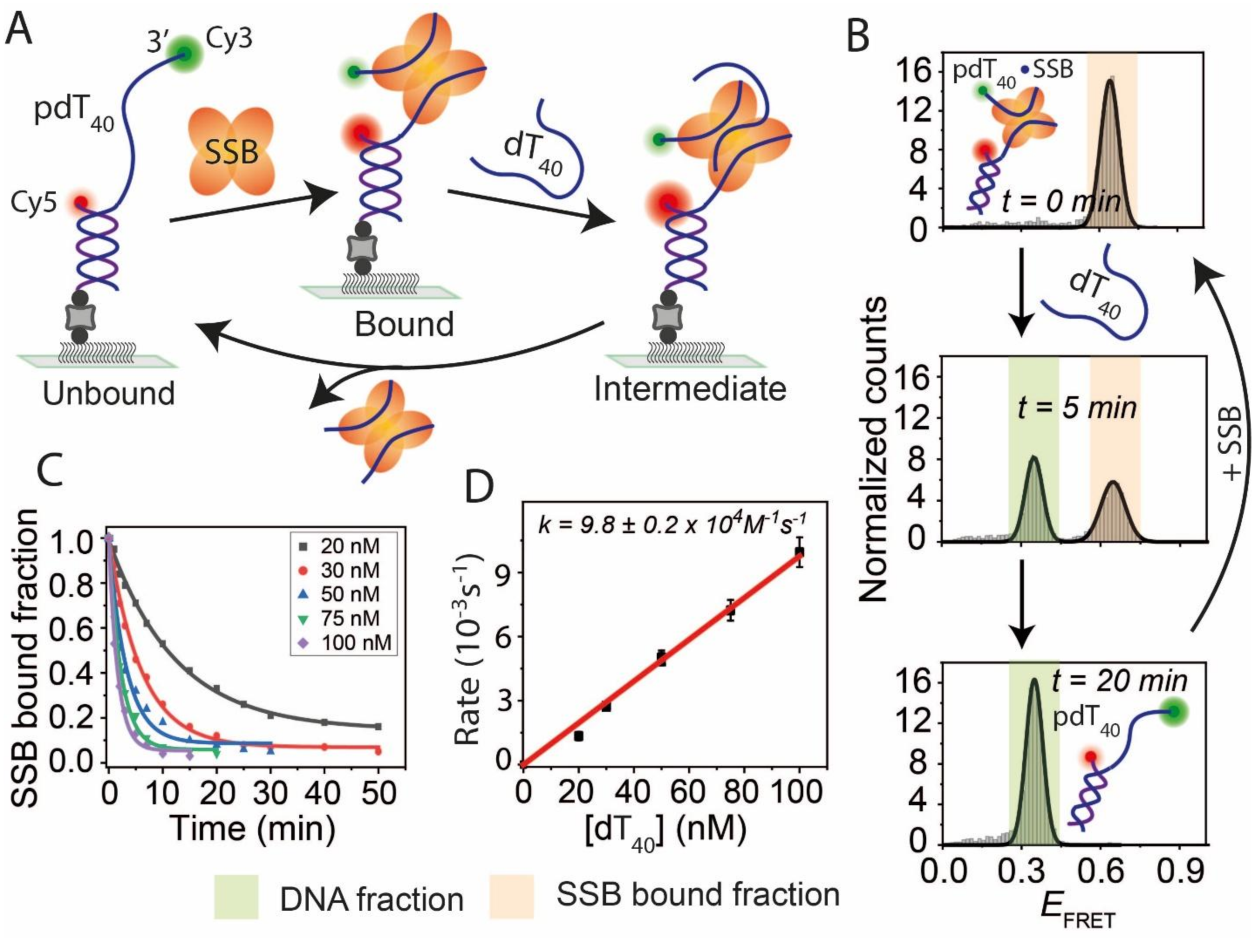


**Supplementary Figure 1:** SSB binding and direct transfer kinetics on pdT40. (A) Schematic of smFRET constructs showing a partial DNA duplex with a 40-nt poly-thymine overhang (pdT40). SSB binds to pdT40, followed by strand transfer to competing ssDNA (dT40). (B) FRET histograms of pdT40 before (bottom) and after SSB binding (top), with the middle histogram showing FRET changes during SSB transfer at indicated times. (C) Single-exponential fitting of the SSB-bound fraction at different dT40 concentrations. (D) Linear fit of SSB transfer rates at varying dT40 concentrations.

**Supplementary Figure 2**

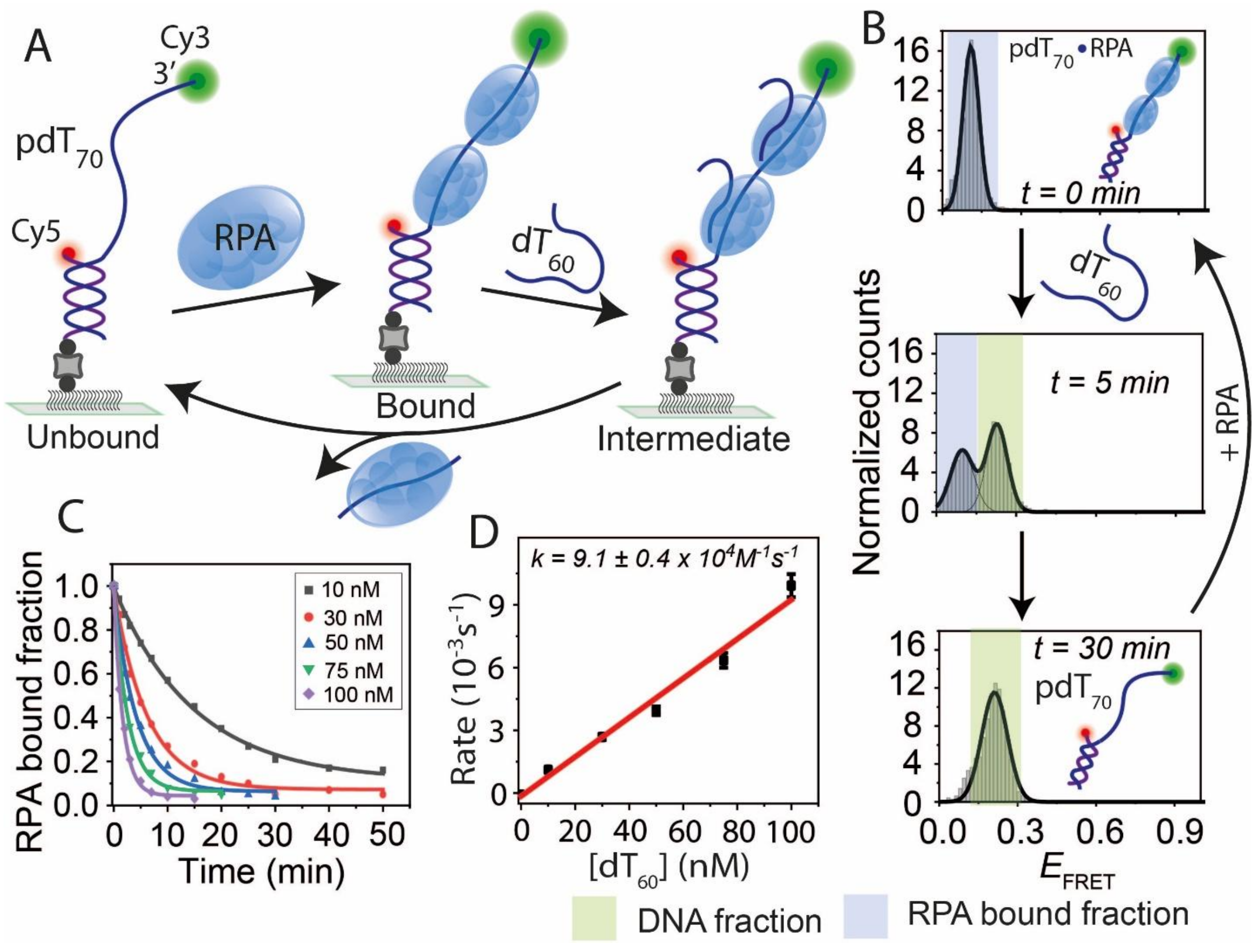


**Supplementary Figure 2:** RPA binding and direct transfer kinetics on pdT70. (A) Schematic of smFRET constructs showing a partial DNA duplex with a 70-nt poly-thymine overhang (pdT70). RPA binds to pdT70, followed by strand transfer to competing ssDNA (dT60). (B) FRET histograms of pdT70 before (bottom) and after RPA binding (top), with the middle histogram showing FRET changes during RPA transfer at indicated times. (C) Single-exponential fitting of the RPA-bound fraction at different dT60 concentrations. (D) Linear fit of RPA transfer rates at varying dT60 concentrations.

**Supplementary Figure 3**

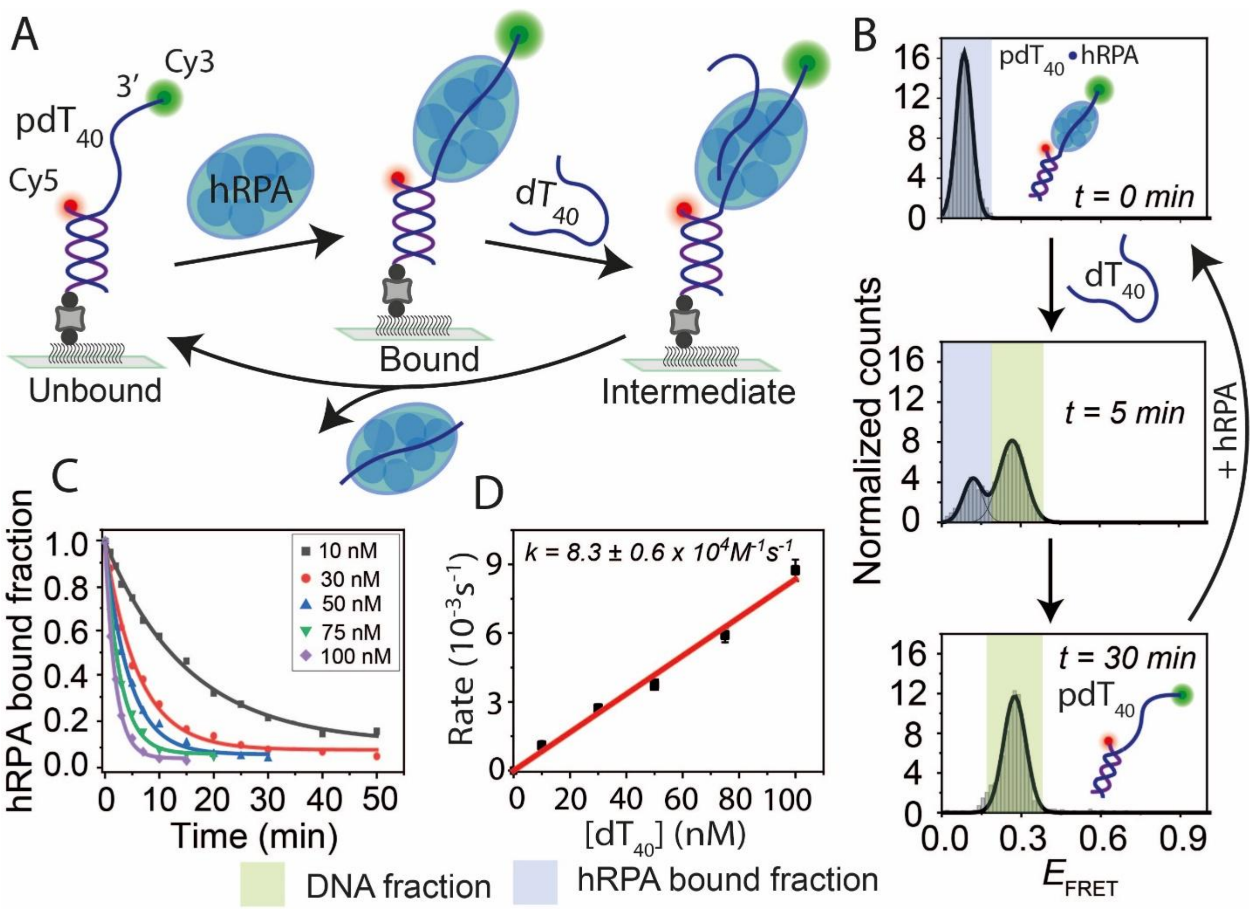


**Supplementary Figure 3:** Human RPA (hRPA) binding and transfer on pdT40. (A) Schematic of smFRET constructs showing a partial DNA duplex with a 40-nt poly-thymine overhang (pdT40). Human RPA (hRPA) binds to pdT40, followed by strand transfer to competing ssDNA (dT40). (B) FRET histograms of pdT40 before (bottom) and after hRPA binding (top), with the middle histogram showing FRET changes during hRPA transfer at indicated times. (C) Single-exponential fitting of the hRPA-bound fraction at different dT40 concentrations. (D) Linear fit of hRPA transfer rates at varying dT40 concentrations.

**Supplementary Figure 4**

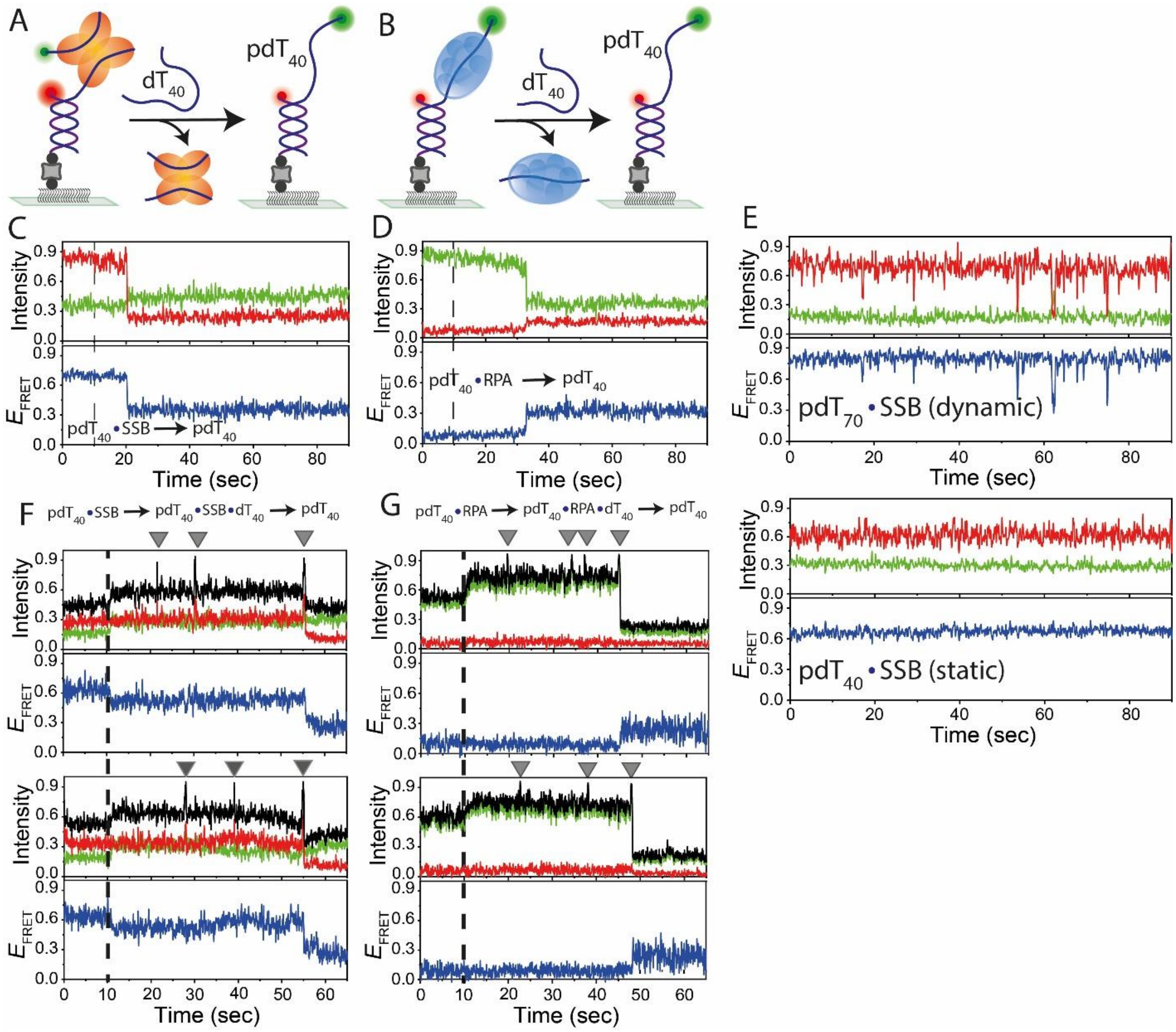


**Supplementary Figure 4:** Real-time smFRET traces of SSB and RPA transfer. (A, B) Schematic smFRET model of pdT70 (pdT40 for RPA) where the bound SSB (A) or RPA (B) transfer to the competing dT60 ssDNA (dT40 for RPA) and regenerate the tethered DNA. (C, D) Representative real-time smFRET traces showing one-step FRET transitions during SSB (C) and RPA (D) transfer to competing ssDNA (100 nM). Dashed lines indicate the addition of competing ssDNA. (E) Representative real-time smFRET traces showing the dynamics of SSB bound to pdT70 (top, dynamic binding) and pdT40 (bottom, static binding). (F, G) Real-time smFRET traces showing SSB (F) and RPA (G) transfer events. Dash line indicate Cy3-ssDNA flow; spikes represent transfer attempts (gray triangles), and the final spike referred to successful transfers with intensity and FRET efficiency changes. The background intensity increases after addition of Cy3-labeled ssDNA. Black, red, green, and blue lines indicate total intensity, Cy5 intensity, Cy3 intensity, and FRET efficiency, respectively.

**Supplementary Figure 5**

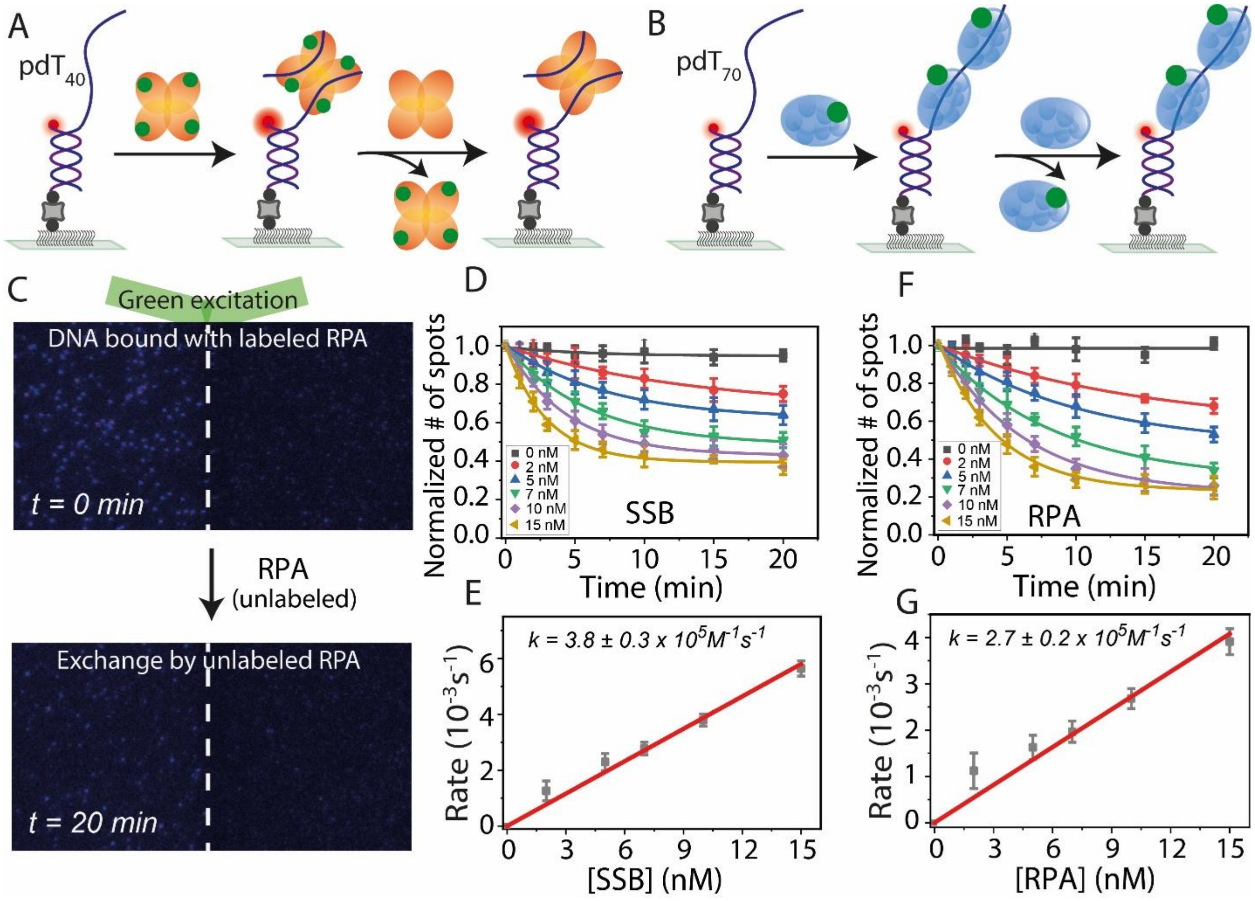


**Supplementary Figure 5:** Protein-protein exchange dynamics of SSB and RPA. (A, B) Schematic representation of SSB (A) and RPA (B) protein-protein exchange, where labeled proteins bound to DNA are replaced by unlabeled proteins. (C) Representative fields of view showing labeled RPA at t=0 min (top) and after exchange with unlabeled protein at t=20 min (bottom) under green laser excitation. (D, F) Single-exponential fits of labeled protein disappearance at different unlabeled protein concentrations for SSB (D) and RPA (F). (E, G) Linear fits of exchange rates at varying protein concentrations for SSB (E) and RPA (G).

**Supplementary Figure 6**

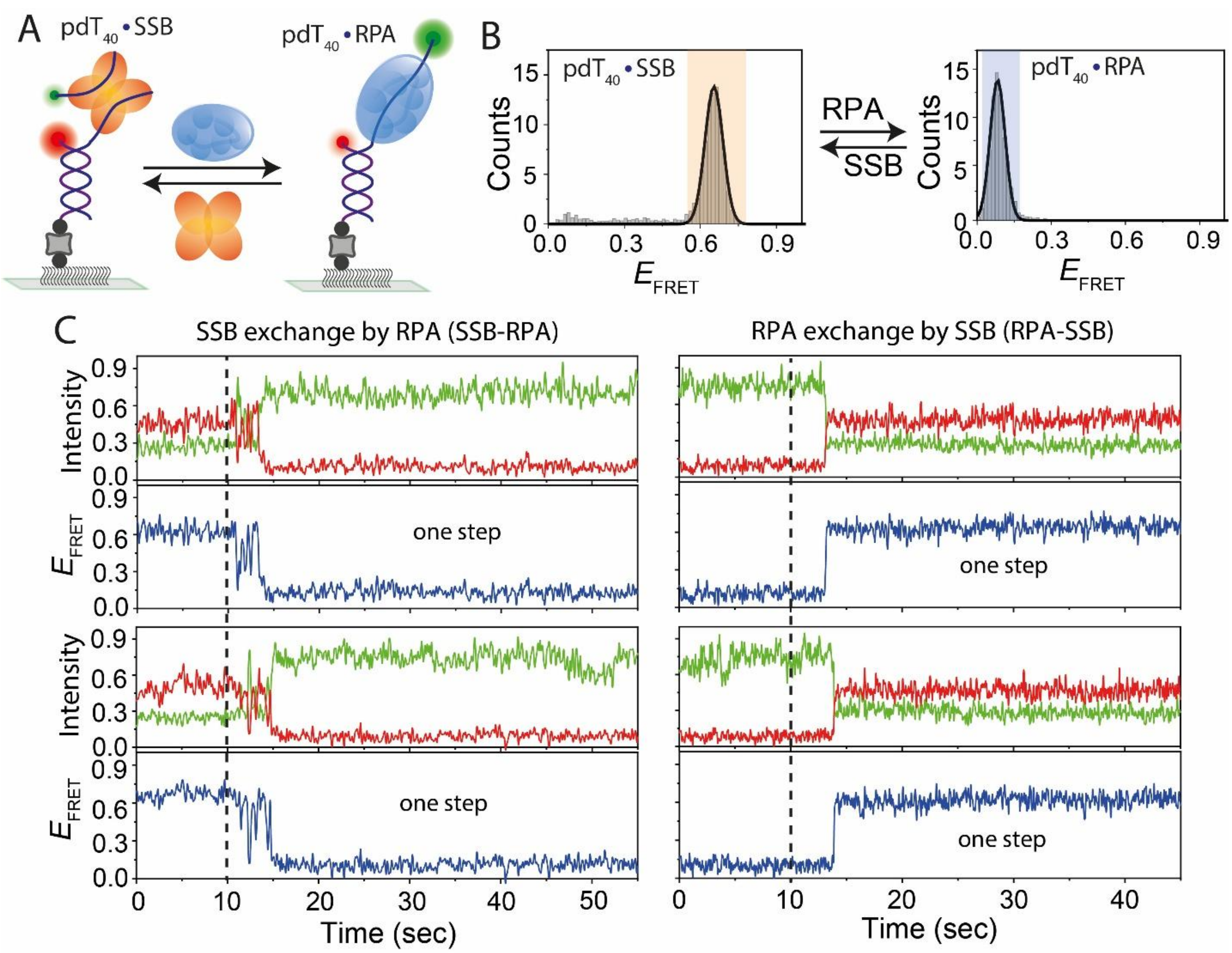


**Supplementary Figure 6:** Hetero-protein exchange dynamics between SSB and RPA. (A) Schematic smFRET model showing hetero-protein exchange between SSB and RPA. (B) FRET histograms of pdT40 showing SSB and RPA binding and their exchange. (C) Real-time smFRET traces showing SSB-to-RPA and RPA-to-SSB exchanges on pdT40. Distinct FRET transitions indicate protein exchange events, with dashed lines marking protein addition.
